# Mesoscale simulations of membrane-tethered reactions to parameterize cell-scale models of signaling

**DOI:** 10.1101/2024.12.04.626844

**Authors:** Kelvin J. Peterson, Boris M. Slepchenko, Leslie M. Loew

## Abstract

Biochemical interactions at membranes are the starting points for cell signaling networks. But bimolecular reaction kinetics are difficult to experimentally measure on 2-dimensional membranes and are usually measured in volumetric *in vitro* assays. Membrane tethering produces confinement and steric effects that will significantly impact binding rates in ways that are not readily estimated from volumetric measurements. Also, there are situations when 2D reactions do not conform to simple mass action kinetics. Here we show how highly coarse-grained molecular simulations using the SpringSaLaD software can be used to estimate membrane-tethered rate constants from experimentally determined volumetric kinetics. The approach is validated using an analytical solution for dimerization of binding sites anchored via stiff linkers. This approach can provide 2-dimensional bimolecular rate constants to parameterize cell-scale models of receptor-mediated signaling. We explore how factors such as molecular reach, steric effects, disordered domains, local concentration and diffusion affect the kinetics of binding. We also develop a general scheme to assess whether simple mass action rate constants can be applied for a given scenario, taking into account the diffusivity of the membrane anchors and tethered binding sites, the initial membrane densities of the reactants and the desired level of completion for the fitted rate constant. We then apply our approach to epidermal growth factor receptor (EGFR) mediated activation of the membrane-bound small GTPase Ras. The analysis reveals how binding of Ras to the allosteric site of SOS, a guanine nucleotide exchange factor (GEF) that is recruited to EGFR, significantly accelerates Ras binding to the SOS catalytic site. A small biochemical network model parametrized with the derived 2D rate constants shows how recruitment of SOS via EGFR can significantly enhance Ras activation.

**SIGNIFICANCE STATEMENT:** In cell signaling, the activation of a surface receptor leads to a cascade of intracellular biochemical events. Many protein interactions occur near the inner plasma membrane surface. However, accurate rate parameters for these steps in models of signaling are rarely available because membrane-tethered reaction kinetics are difficult to experimentally measure. Here, we use a highly coarse-grained molecular simulator to model the kinetics of reactions between binding sites that are tethered to a membrane. We can fit these simulation outputs with 2-dimensional rate laws to obtain rate constants that can be used to build complex models of cell signaling. The derived rate constants can also be analyzed to understand the key biophysical features controlling the kinetics of bimolecular membrane reactions.

## Introduction

The cell membrane responds to and integrates electrical, mechanical and chemical signals from the extracellular environment. For chemical signals, the initial step is binding of a ligand to an external binding site on a membrane receptor protein. This triggers a chain of events that typically involves a change of state of the cytoplasmic receptor domain and subsequent recruitment of adapter proteins, enzymes and/or cytoskeletal regulators to further evoke a cell biological response. Mathematical modeling of signaling pathways is a powerful tool to systematically organize the experimental knowledge we have about these complex systems and then develop predictions, through simulations, to inspire new experiments (1-3).

A challenge in developing cell signaling models is the acquisition of the appropriate experimentally-grounded input parameters. Often, kinetic data is available from *in vitro* biochemistry and this has served the mature field of metabolic modeling very well. However, rate parameters are less available and more difficult to measure for signaling pathways and networks. Among the key challenges is that many of the essential steps are associated with the plasma membrane, where multiple molecules are recruited before a messenger ultimately diffuses to an intracellular target (e.g. the nucleus). It is experimentally difficult to measure reaction rates on membranes because complex reconstituted bilayer methods must be used (4), so available data is commonly derived from volumetric measurements. While such quantitative data is useful, it can be challenging to translate rate parameters derived from 3D solution to the very different biophysical environment of a 2D membrane.

Indeed, the biophysics of membrane associated reactions has a long scientific history. An early focus of investigation, initiated with a classic paper by Adam and Delbruck (5), was the difference between 2D and 3D diffusion limited reactions; they argued that the 2-step process of absorbing a cytosolic molecule to the membrane and subsequent 2D search for an enzyme or binding partner might offer a kinetic advantage over a fully 3D search. This argument emerges from the fundamental difference between 2D and 3D diffusion: a diffusing molecule in 2D is guaranteed to eventually find its target, while no such guarantee can be made in 3D (6). In practice, Adam and Delbruck’s assertion that reactions in membranes have an inherent advantage has been disputed using subsequent experimentally determined realistic parameters and biological scenarios; this is because of the generally low diffusivity of membrane embedded molecules compared to the same sized molecule diffusing in the cytosol (7).

However, regardless of whether 2D reactions are advantageous over 3D, a consequence of the Adam and Delbruck (5) analysis is that there is a fundamental difference between 2D and 3D bimolecular reactions. Reactions in 3D can generally be described by mass action kinetics, where rate constants are truly constant, i.e independent of time and reactant concentrations. However, bimolecular reactions on a surface cannot be fundamentally described by mass action kinetics (8-14). With regard to cell signaling, theoretical analyses of reactions at membranes have been extended and elaborated to consider both diffusion-limited and reaction-limited bimolecular kinetics (1,8,10,11). Intuitively, for reaction-limited cases (where diffusion is fast compared to the intrinsic reaction rate upon encounter) the Adam and Delbruck idea is not pertinent and the reaction can be well described by simple mass action kinetics. In particular, Yogurtcu and Johnson (10) developed the theory to determine where the transition from reaction-limited to diffusion limited 2D kinetics occurs – i.e. when mass action in 2D is an adequate description or when a more complex time-dependent parameterization of the rate is required.

These pioneering studies treated membrane reactions as strictly two-dimensional surface events. However, most biological membrane associated reactions actually occur in the immediately adjacent cytosol, with interacting sites tethered to the membrane through lipid or protein anchors. A well-known feature of anchoring bimolecular reactions to a membrane is the effect of locally increased effective concentration: compared to the same reaction by the same number of molecules within the cell volume, anchoring the reaction to the membrane generally increases concentration by confining the reaction volume to a thin layer above the membrane (15). The thickness of that layer is often parametrized as *h*, sometimes called the “confinement length” (16,17). Roughly, *h* is related to the distance the binding sites can sample above the membrane surface. The smaller *h*, the greater the effective concentrations of binding sites and the faster the membrane associated bimolecular kinetics. In principle, *h*, can be determined by analyzing detailed molecular dynamics simulations on the flexibility and motions of binding domains tethered to the membrane (17,18). Recently, the concept of *molecular reach* was introduced as a more general framework for assessing how molecular structure influences the steady state phosphorylated fraction of a membrane bound substrate interacting with a tethered kinase (19). The *reach* is defined as the distance of the kinase site from the membrane anchor and is directly related to *h* when the anchor diffuses freely in the membrane. However, when lateral diffusion is restricted (e.g. within large signaling clusters such as the immune synapse), binding sites with longer *reach* may have an advantage in being able to find more binding partners; for such diffusion-limited scenarios, this increase in reach can outweigh the decrease in local concentration associated with increased *h* (19).

Thus, it is clear from these foundational studies that the membrane environment and the structural features of interacting membrane-bound molecules must be considered when converting a measured 3D on-rate to a 2D on-rate suitable for cell-scale continuum models based on ordinary or partial differential equations (ODEs or PDEs). In this work, we show how this can be done using simulations from SpringSaLaD (20) to derive 2D rate constants. Importantly, we are treating tethered binding sites. This is in fact the more general situation in cell signaling where a more-or-less flexible protein domain containing the binding site extends into the cytosol a significant distance from the actual membrane – typically much longer than the actual thickness of the lipid bilayer. Because our method accounts for the structure of the protein (albeit very coarsely), the distance of the binding sites from the membrane as well as other biophysical features are accounted for in the estimation of the 2D on-rate constant.

SpringSaLaD uses a series of variously sized spherical sites linked together with stiff springs to coarsely model the key structural features of macromolecules such as flexibility, excluded volume and binding site localization. Each sphere within the molecule can be assigned its own diffusion coefficient and Brownian diffusion is simulated via a Langevin dynamics algorithm. The molecule can be tethered to a surface, representing a membrane, via a specialized anchoring sphere that has a lateral diffusion coefficient; the rest of the molecule, including the spheres designated as binding sites, are free to explore the volume above the membrane within their reach. Naturally therefore (and particularly advantageous for the purpose of this work), volumetric rate constants are used for bimolecular rate expressions even for membrane-bound molecules. In previous work, this feature was used to show how multivalent clustering is enhanced when one of the interacting molecules is tethered to a membrane (21).

To determine 2D rate constants, we fit the stochastic kinetics simulated with SpringSaLaD to a deterministic mass-action rate law based on the corresponding surface densities of the binding partners. Using idealized structures, we validate this procedure against theoretical calculations. We then explore how molecular structural features (e.g. tether length and stiffness, steric access), surface density and lateral diffusion affect binding kinetics. Additionally, we develop a theory for membrane-tethered reactions, similar to that of Yogurtcu and Johnson for purely 2D reactions (10), to estimate the applicability of the mass action rate law for varying tether lengths, anchor diffusion coefficients, reaction rates, initial surface densities and the desired level of completion of the reaction. Then, as a biologically relevant example, we apply this approach to recruitment of SOS to the epidermal growth factor receptor (EGFR). SOS is the G-protein exchange factor (GEF) for Ras (22) and a better understanding of SOS activation of Ras emerges from this analysis. Thus, we offer a procedure for parameterizing membrane kinetics that serves to bridge molecular to cell scale simulations of cell signaling.

## Methods

All simulations were performed with SpringSaLaD v. 2.3.4 (https://vcell.org/ssalad). While the full physics and math behind the calculations performed by SpringSaLaD can be found in the original publication (20), we provide a brief overview of its functionality. A molecule in SpringSaLaD is approximated as a series of variably sized hard spheres, representing protein domains, connected by stiff springs. Random forces (chosen from a normal distribution with a variance determined from the diffusion coefficient assigned to each sphere by the user) impinge on each sphere at each time step and are transmitted to neighboring spheres through the stiff springs. The software models membranes as a planar surface within the volume of the 3D simulation domain. The membrane can have embedded spheres that serve as diffusing anchors (with user-assigned 2D diffusion coefficients) linked to spheres that extend into the volumetric space. Each sphere may also be designated as the site of a reaction. The user can input macroscopic rate constants for reactions and the software converts these to reaction probabilities within each time step; for bimolecular reactions, this conversion also depends on the radii of the spherical binding sites and the sum of their diffusion coefficients. SpringSaLaD allows users to manually edit molecules to fully customize structure and molecule flexibility. Orientational approaches between spherical binding sites can be restricted by the appropriate placement of neighboring spheres adjacent to the respective binding sites within the reacting molecules. To build coarse grained molecular structures used in SpringSaLaD for Fig. 3, we used AlphaFold2 (23,24) to generate PDB file estimates of protein structures for EGFR, Grb2, SOS, and Ras via input of entire amino acid sequences. These PDB files are converted to highly coarse-grained molecular models via the mol2sphere (25) utility embedded in SpringSaLaD. In some cases, we manually edited the structures to capture their essential features from measurements on the PDB structures, as visualized in PyMol (Schrödinger, Inc.). In particular, for SOS we subdivide the CDC25 and REM domains into multiple smaller spherical sites with only two sites (the allosteric and catalytic sites for RAS) capable of participating in a binding reaction. This process maintains the structural characteristics of these domains, while ensuring that the binding radius of the domain is not artificially inflated. Modeling disordered regions, such as the PRM region of SOS, can be challenging due to low confidence in the AlphaFold2-generated geometry of these regions. To model disordered domains, we use PyMOL to measure the length of entire straight chain amino acid sequences, then model this sequence in SpringSaLaD using 1.0 nm diameter sites connected by 3.1 nm linkers (in SpringSaLaD, a string of relatively small spheres connected by links that are larger than the sphere diameters can be used to model flexible (i.e. disordered) domains). Binding reactions in all simulations have rates input in terms of µM^−1^s^−1^; in our simulations, successful binding results in 1nm (Tables 1 and 2, Fig. 4) distances between the surfaces of the spherical sites. The default simulation time step of 10ns was sufficiently accurate for all the SpringSaLaD simulations in this work (as checked by testing 2 ns timesteps and noting no significant differences). Simulations for Figs. 1 and 2 were run with 40 molecules, while Figure 4 used 200 molecules of Ras and 20 complexes of EGFR-Grb2-SOS to proportionally reflect the difference in concentrations between Ras and SOS molecules *in-vivo* (26,27). The 3D problem domain had a z coordinate of 100nm, with the membrane placed in the XY plane at Z=10nm; the boundaries at the edges of the domain have reflective boundary conditions. X and Y coordinates were set to provide the desired initial surface densities. Initial placement of the molecules was random and different for each trajectory. A set of 100 trajectories are simulated in parallel using the Center for Cell Analysis and Modeling High Performance Compute Cluster (https://health.uconn.edu/high-performance-computing/resources/). One run for Tables 1 and 2 required approximately 3 hours and one run for Fig. 4 required approximately 30 hours. SpringSaLaD input files are in Supporting Information and provide all the geometric details for the molecules in each computational experiment.

**Table 1.** Results for single stiff link, binding site*D*_*vol*_=1.0µm^2^/s, *h*=0.0055µm 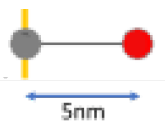.

| $k_{on}^{(vol)}$<br>($\mu\text{M}^{-1}\text{s}^{-1}$) | Surface density<br>(molecules/$\mu\text{m}^2$) | $D_{mem}$<br>($\mu\text{m}^2/\text{s}$) | $k_{on}^{(h)} = k_{on}^{(vol)}/h/602.2$<br>($\mu\text{m}^2\text{molecules}^{-1}\text{s}^{-1}$) | $k_{on}^{(mem)}$, SpringSaLaD fit<br>($\mu\text{m}^2\text{molecules}^{-1}\text{s}^{-1}$) | Relative<br>Sq Dev | $\delta$ |
| --- | --- | --- | --- | --- | --- | --- |
| 0.01 | 2500 | 0.01 | $3.0 \times 10^{-3}$ | $2.9 \times 10^{-3}$ | $3.0 \times 10^{-5}$ | 0.09 |
| 0.01 | 2500 | 1.0 | $3.0 \times 10^{-3}$ | $3.1 \times 10^{-3}$ | $3.1 \times 10^{-5}$ | 0.001 |
| 0.01 | 25 | 0.01 | $3.0 \times 10^{-3}$ | $2.4 \times 10^{-3}$ | $2.9 \times 10^{-5}$ | 0.13 |
| 0.01 | 25 | 1.0 | $3.0 \times 10^{-3}$ | $3.0 \times 10^{-3}$ | $2.3 \times 10^{-5}$ | 0.002 |
| 1.0 | 2500 | 0.01 | 0.30 | 0.08 | $7.4 \times 10^{-3}$ | 0.78 |
| 1.0 | 2500 | 1.0 | 0.30 | 0.26 | $2.6 \times 10^{-4}$ | 0.04 |
| 1.0 | 25 | 0.01 | 0.30 | 0.02 | $1.5 \times 10^{-3}$ | 0.9 |
| 1.0 | 25 | 1.0 | 0.30 | 0.20 | $4.9 \times 10^{-4}$ | 0.09 |
| 1.0<br>($k_{off}=10^3\text{s}^{-1}$) | 2500 | 0.01 | 0.30 | 0.18 | $1.3 \times 10^{-3}$ | |
| 1.0<br>($k_{off}=10\text{s}^{-1}$) | 25 | 0.01 | 0.30 | 0.1 | $6.2 \times 10^{-2}$ | |

**Table 2.**
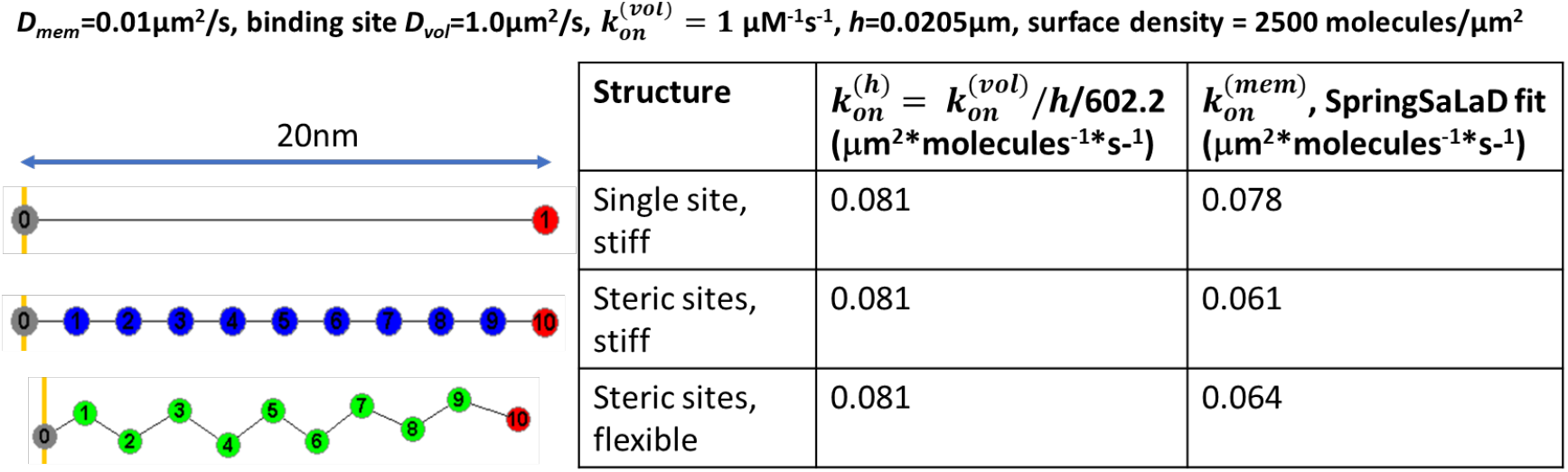
Dimerization of idealized structure types.

| Structure | $k_{on}^{(h)} = k_{on}^{(vol)}/h/602.2$<br>($\mu\text{m}^2*\text{molecules}^{-1}*\text{s}^{-1}$) | $k_{on}^{(mem)}$, SpringSaLaD fit<br>($\mu\text{m}^2*\text{molecules}^{-1}*\text{s}^{-1}$) |
| --- | --- | --- |
| Single site,<br>stiff | 0.081 | 0.078 |
| Steric sites,<br>stiff | 0.081 | 0.061 |
| Steric sites,<br>flexible | 0.081 | 0.064 |

**Figure 1.**
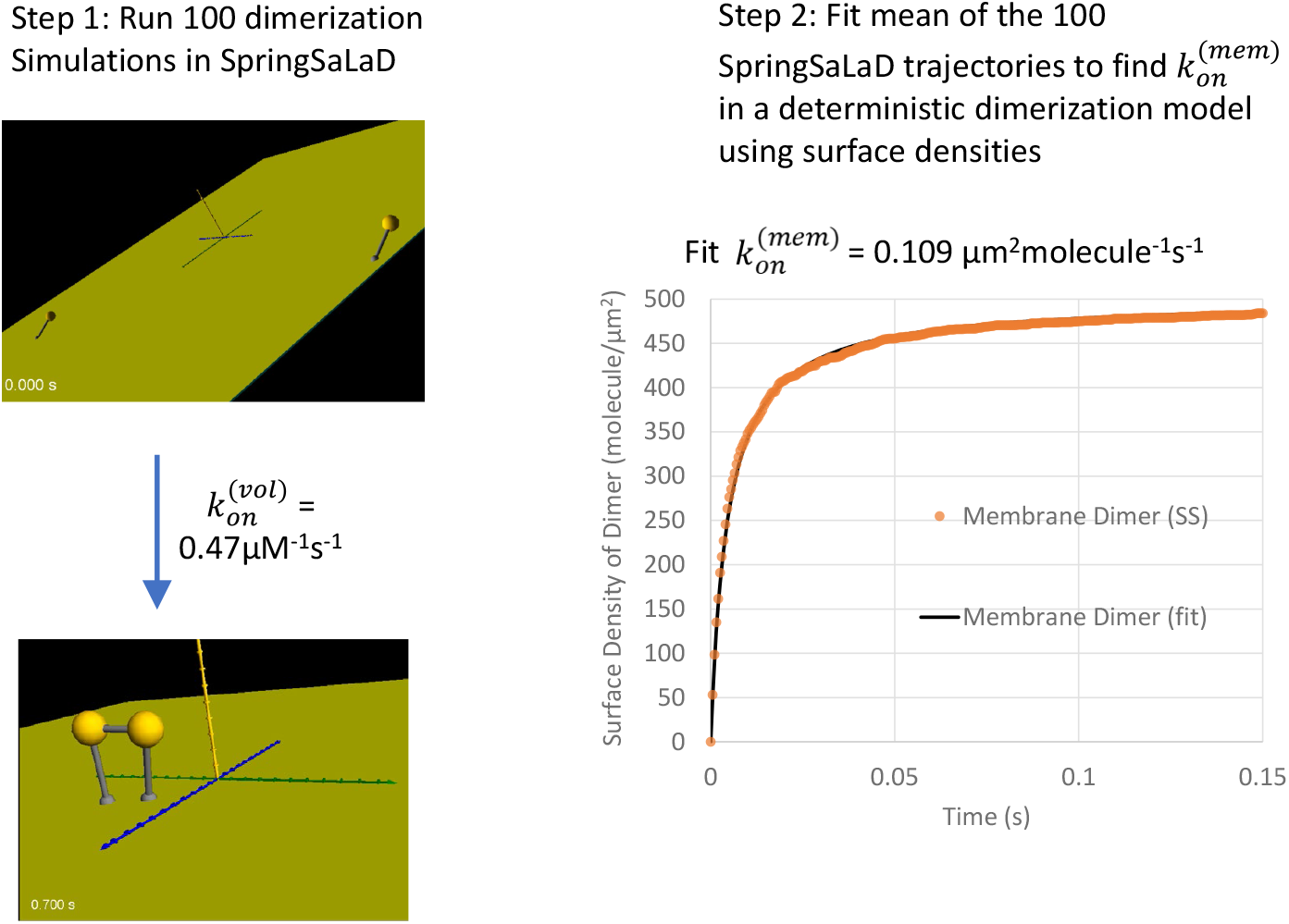
Workflow for finding membrane bimolecular binding rate constants, 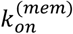, in terms of surface densities. Step 1 is to run 100 SpringSaLaD trajectories with the coarse-grained model of the binding partners. Illustrated are a pair of simple monomers (top) consisting of 2nm diameter spheres (yellow) tethered to a 1nm membrane anchor sphere (gray) with a 5nm link; both the anchor and binding spheres are assigned the diffusion coefficient of 1 µm^2^/s. At the bottom, the product dimer is depicted. Step 2 consists of fitting the average of 100 outputs from stochastic SpringSaLaD (SS) simulations with a deterministic (ODE) non-spatial model of surface-bound dimerization.

**Figure 2.**
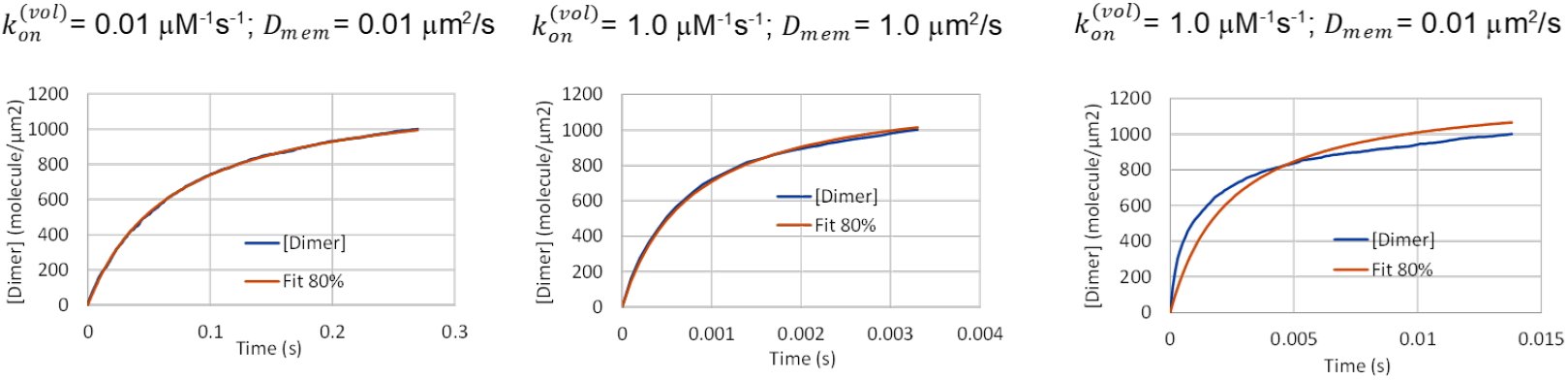
Examples of SpringSaLaD simulation data and their fit to surface-confined mass action dimerization kinetics. The initial surface density of monomers in each case is 2500 molecule/µm^2^. The diffusion coefficient of the anchor and the volumetric binding rate constant of the binding sites are indicated above each graph, corresponding respectively to rows 1, 6 and 5 of Table 1. Each of the SpringSaLaD simulations were run to 80% completion.

**Figure 3.**
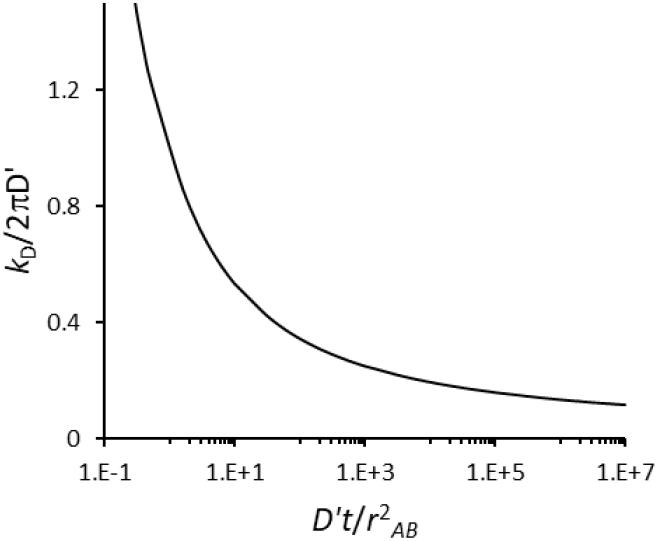
Time dependence of *k*_*D*_ in 2D.

**Figure 4.**
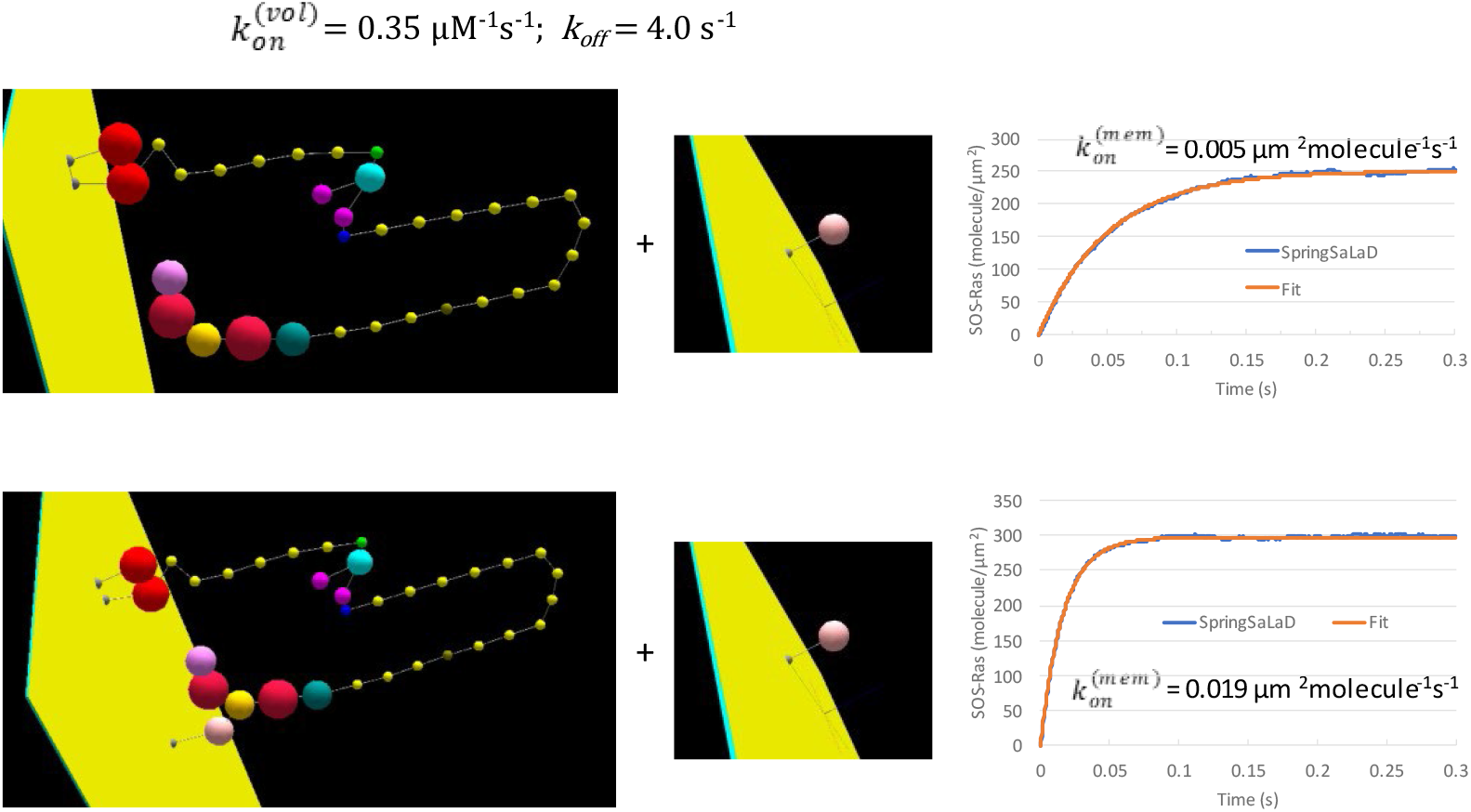
Membrane binding of Ras to the catalytic site of Receptor-bound SOS. Top: direct binding. Bottom: after pre-binding of Ras to the allosteric site. The molecular structures are approximated by using the SpringSaLaD 3D editing utility based on atomic structures derived from AlphaFold2. The top left structure is an EGFR cytoplasmic domain anchored to the membrane (red kinase domain, followed by yellow disordered tail capped by a phosphotyrosine in green); the latter is linked to a cyan SH2 domain in Grb2; one of its magenta SH3 domains is linked to an olive PRM on the end of the disordered region of SOS; the violet SOS binding site for the catalytic domain of Ras is indicated with an asterisk (*). The bottom left structure is identical, except that the pink allosteric site on SOS is prebound to Ras. The Ras structures are shown in the center with the yellow binding sites indicated by an asterisk. The input rate constants for the SpringSaLaD simulations are shown at the top, corresponding to the volumetric on rate for Ras binding to the catalytic site of SOS. The EGFR anchor diffusion coefficient is 0.01µm^2^/s. All other site diffusion coefficients are 1.0µm^2^/s. For each condition, 20 EGFR-Grb2-SOS molecules react with 200 Ras molecules on a 250nmX250nm membrane surface to generate 100 SpringSaLaD trajectories. Their means were fitted to a deterministic 2D rate law to derive 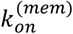, fixing *k*_*off*_ at 4.0 s^−1^; results for the 2 conditions are shown on the right.

The mean time dependence of dimer surface density [*dimer*]_*t*_, based on 100 SpringSaLaD runs for each of the parameter sets in Tables 1 and 2 and Fig. 4, were fitted with a deterministic 2D mass action rate law to obtain 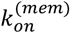. For irrreversible dimerization (Table 1, first 8 rows and Table 2), a fit to the analytical mass action expression for the appearance of dimer (Eq. 1) used the Excel solver.

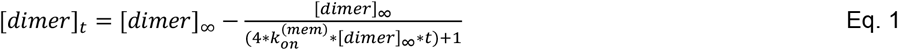

For reversible dimerization (Table 1, rows 9 and 10) and for the fits in Fig. 4, we used the COPASI (28) parameter estimation tool within Virtual Cell (VCell) (29,30). The latter can be accessed in the VCell published BioModel “Peterson Figure 4: Ras-SOS_Binding_fit_to_SpringSaLaD”. All these results with some further analysis can also be found in the spreadsheets included in the Supporting Information. The VCell model related to Fig. 5 can be found in the VCell database with the name “SOS_Recruitment_Ras_Binding”

**Figure 5.**
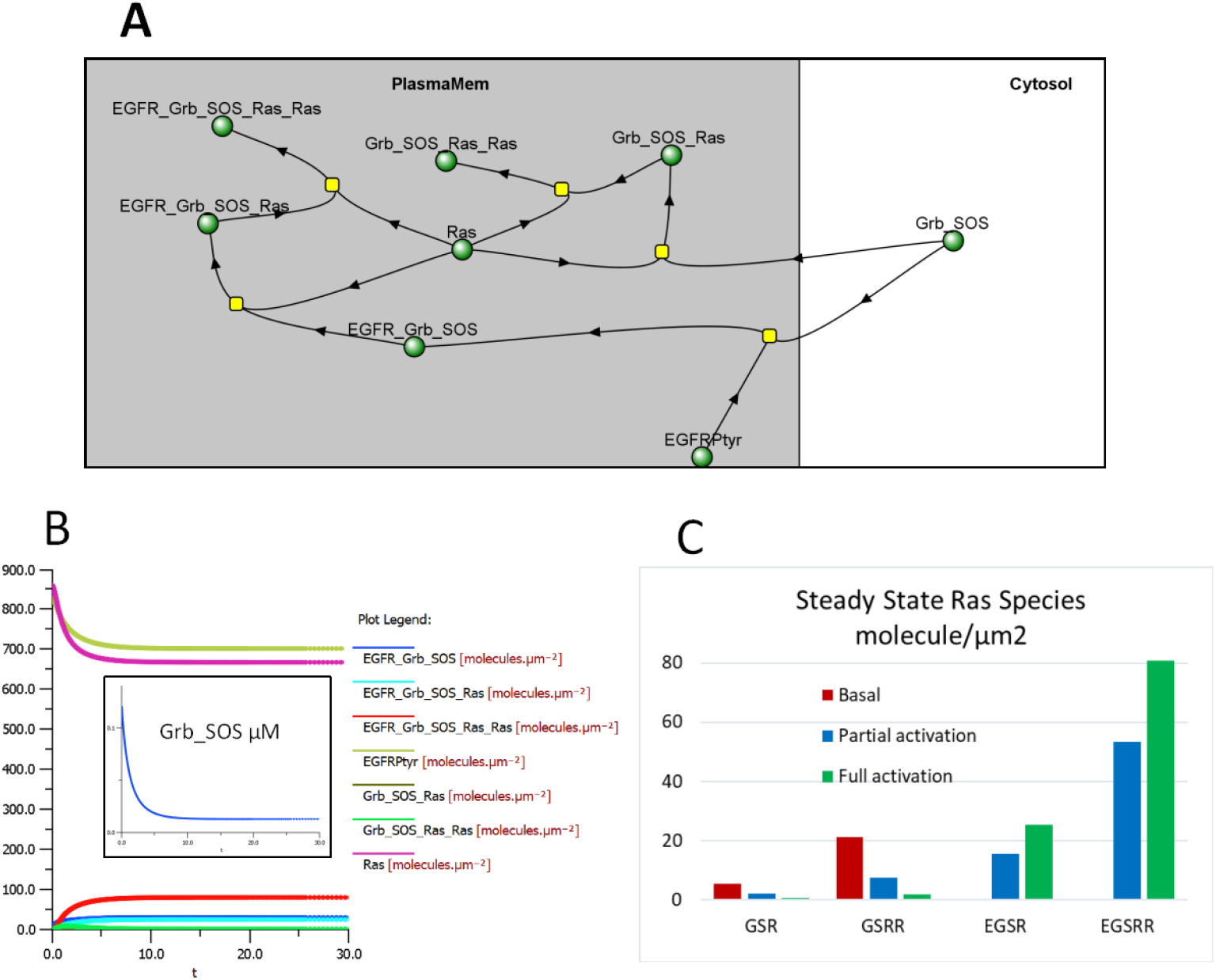
A simple VCell model and simulation results for EGFR signaling to Ras. A. Reaction network identifying all the variables (see also the corresponding reactions in Table 3.). B. The kinetics reach steady state within 30s. The primary plot shows all the membrane species and the inset shows Grb_SOS in the cytosol. Steady state surface densities of the 4 different species of 1 or 2 Ras bound to Grb_SOS or EGFRPtyr_Grb_SOS under 3 conditions: “Basal”, “Partial activation” and “Full activation” corresponding respectively to 0, 200 and 840 molecules/µm^2^ initial EGFRPtyr.

## Results

### The general approach.

For bimolecular reactions, SpringSaLaD determines the microscopic probability of 2 binding sites forming a bond as they diffuse within a reaction radius that is slightly larger than the sum of their physical radii. The input to the algorithm is simply the macroscopic volumetric on rate constant (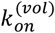, units of µM^−1^s^−1^) and the diffusion coefficient of the individual spheres. Full details on the derivation of the reaction probability and a thorough validation of its accuracy can be found in the original paper describing SpringSaLaD (20). Importantly for the purposes of this work, the rate of binding for sites that happen to be tethered to a membrane are still treated as volumetric, because the spherical sites are located in the volume compartment even while they are constrained with links to the 2D membrane surface.

**Table 3.** Rate constants for the VCell model.

| Reaction | $k_{on}$ | $k_{off}$ | Source |
| --- | --- | --- | --- |
| EGFR<sub>Ptyr</sub> + Grb_SOS → EGFR_Grb_SOS | $1.0 \mu\text{M}^{-1}\text{s}^{-1}$ | $0.3 \text{ s}^{-1}$ | (42) |
| Grb_SOS + Ras → Grb_SOS_Ras | $0.27 \mu\text{M}^{-1}\text{s}^{-1}$ | $4.0 \text{ s}^{-1}$ | (22) |
| EGFR_Grb_SOS + Ras → EGFR_Grb_SOS_Ras | $0.005 \mu\text{m}^2\text{molecule}^{-1}\text{s}^{-1}$ | $4.0 \text{ s}^{-1}$ | (22) with Fig. 4 |
| Ras + EGFR_Grb_SOS_Ras → EGFR_Grb_SOS_Ras_Ras | $0.019 \mu\text{m}^2\text{molecule}^{-1}\text{s}^{-1}$ | $4.0 \text{ s}^{-1}$ | (22) with Fig. 4 |
| Ras + Grb_SOS_Ras → Grb_SOS_Ras_Ras | $0.019 \mu\text{m}^2\text{molecule}^{-1}\text{s}^{-1}$ | $4.0 \text{ s}^{-1}$ | (22) with Fig. 4 |

Figure 1 illustrates this for a simple dimerization reaction where the two yellow binding sites are tethered to the membrane anchor (gray sphere) by a 5nm link; in these simulations both the anchor and tether sites are given identical diffusion coefficients of 1µm^2^/s. We ran 100 SpringSaLaD simulations each with 40 dimerizing molecules (Fig. 1 only shows two molecules for clarity). The mean trajectories for these 100 runs are then fitted with a deterministic mass action membrane binding model in terms of surface densities using either COPASI or Virtual Cell (although an analytical solution can be also fit for simple dimerization) (Step 2 in Fig. 1). The output binding rate constant, 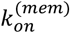 in units of µm^2^molecule^−1^s^−1^, can then be used to parameterize larger deterministic or stochastic models (that utilize mass action rate constants as inputs) with molecule numbers (>1000) or timescales (>10 s) that would be too large for even highly coarse-grained molecular simulators like SpringSaLaD. Also, we emphasize that binding rates are typically determined experimentally using in vitro volumetric measurements; that SpringSaLaD uses volumetric rate constant inputs, makes the procedure in Fig. 1 especially appropriate and convenient. The results in Fig. 1 show that a 2nm diameter binding site with an on rate for dimerization of 0.47 µM^−1^s^−1^ and tethered to a membrane surface through a 5nm link, can be modeled as a 2D surface reaction with an on rate, 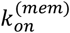, of 0.109 µm^2^molecule^−1^s^−1^. Using idealized models, we now explore how various structural and biophysical parameters control 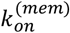 and when mass action rate constants may not be appropriate to describe membrane-associated binding kinetics.

### Effects of volumetric on-rate, 2D diffusion rate, surface density and structural features on dimerization kinetics in idealized systems

We start by analyzing the case of a 5nm stiff tether between a 1nm diameter binding sphere and the membrane. The diffusion coefficient for the binder sphere is set to 1µm^2^/s. Table 1 gives results for all combinations of two volumetric on rate constants, two surface densities and two membrane diffusion coefficients. All the dimerization rate laws are irreversible except for the last two rows, where the off-rate constant is indicated. The 2D on-rate constant derived by fitting the SpringSaLaD simulation, 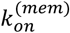, is in the fifth column; for consistency, all these fits are performed for kinetics at 80% completion (the impact of the choice of the level of completion is discussed in the next section). For comparison we also provide the 2D on-rate constant, 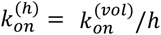, where *h* is the linker length plus the radius of the binder sphere. If *h* is in units of µm, dividing by a unit conversion factor of 602.2 converts 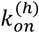 from units of µM^−1^s^−1^ to units of µm^2^molecules^−1^ s^−1^; 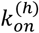 is an equivalent 2D binding rate constant of freely diffusing monomers confined within a thin volume with a height *h* adjacent to the membrane. The Relative Sq Dev column provides the ratio of the sum of the squared deviations to the sum of the squared SpringSaLaD mean values; this ratio provides a measure of the goodness of fit to the bimolecular mass action rate law, with anything less than ~10^−3^ representing a good fit. Examples of the fits are shown in Figure 2. The last column, δ, is a measure of how applicable mass action kinetics are for these combinations of parameters; this will be further discussed in the next Section.

We looked at two anchor diffusion coefficients corresponding to that of a large transmembrane protein domain (*D*_*mem*_= 0.01 µm^2^/s) and a lipid anchor (*D*_*mem*_=1 µm^2^/s). The first row of Table 1 considers a case where diffusion of the binder (*D*_*vol*_=1 µm^2^/s) is 100 times that of the anchor site in the membrane. Because the linker is stiff, the anchor acts as a pivot and the binder rapidly moves within a hemispherical shell to effectively create a reaction region with a thickness slightly larger than the 1-nm diameter of the binder sphere. Because the region of spatial overlap of the two shells where the binding may occur is restricted, it might seem surprising that 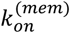 is so closely approximated by 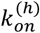. However, as shown in the Supporting Text I, the reaction probability is enhanced because of the effectively higher density of binding sites within this restricted region, which compensates for the smaller spatial overlap. Thus, the calculations in the Supporting Text I both explain and validate the SpringSaLaD results. The second row of Table 1 corresponds to the case where the anchor and the binder have the same fast diffusion. In this scenario, that 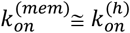 is intuitive, because the effect of the tether in this case essentially reduces to confining the binders within the layer adjacent to the membrane. The solutions in the Supporting Information assume that the molecular distributions are spatially uniform at any time (i.e., well-mixed), which pertains best to the cases where the reaction is slow on the timescale of <u>anchor</u> diffusion (reaction-limited kinetics).

It has been shown that the rates of membrane reactions may be susceptible to deviations from a simple mass action rate law (8,10,14,31-35), which manifests themselves as significant changes of the apparent rate constant with initial surface density. These changes need to be assessed before using the rate constants we obtain by the procedure of Fig. 1 in large cell-scale models. We start by a quantitative exploration of how variations of some of the key parameters in the test cases of Table 1 might expose deviations from mass action. We probed for this by decreasing the initial density by a factor of 100. The combinations of *D*_*mem*_ and 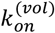 in the third and fourth rows of Table 1 resulted in relatively small changes in 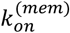, indicating that the mass action rate law applies to these cases. We further tested this by increasing the 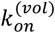 by a factor of 100 in the lower half of Table 1. Clearly, for the case of *D*_*mem*_= 0.01 µm^2^/s, there is a strong dependence on surface density. Furthermore, the system under these conditions develops depletion zones, as discussed below. Thus, the combination of 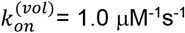 and *D*_*mem*_= 0.01 μm^2^/s present cases where mass action rate laws would not apply. Of course, the appropriateness of the mass action rate law can also be judged by the goodness of fit when determining 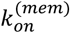 from the SpringSaLaD simulation (last column of Table 1); as demonstrated in Fig. 2, the case of 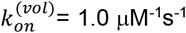 and *D*_*mem*_ = 0.01 μm^2^/s is not approximated very well by a mass-action dimerization rate. The simulation results are initially faster and then ultimately slower than the best fit that assumes mass action kinetics. This is because the monomers whose binding sites are initially close to each other (effectively within the “reach” of the tether) will react, but leave behind depletion zones where monomers are too far away from potential binding partners (33,34). These isolated monomers can be discerned toward the end of Movie 1, which presents an example trajectory for this case. A discussion of how to more generally assess the appropriateness of mass action kinetics is provided in the next Section.

Consistent with the idea of depletion zones, the last 2 rows of Table 1 show that when reversibility is introduced, the fitted on-rate constant increases (compare, respectively, to rows 5 and 7 of Table 1). This is because when dimerization is reversible, free monomers can reappear to fill in depletion zones, thereby countering the slow diffusion. While the mass action kinetics approximations are still poor, especially for the low density case, the respective mean steady-state densities of dimers in the SpringSaLaD simulations, 570 and 5.7 molecules/µm^2^, are consistent with the *thermodynamic* law of mass action (9). According to thermodynamics, the ratio of the squared monomer surface density and the dimer surface density is equal at steady state to the equilibrium dissociation constant *K*_*d*_, which is determined by the strength of the interaction between the monomers and thus is independent of reaction kinetics. When kinetics obey mass action, *K*_*d*_ is equal to the ratio of off and on rate constants. In our case, 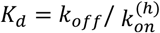, because the interactions of the binding sites in the 3D volume above the membrane obey mass-action kinetics and 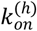 is the 3D association rate constant adjusted for the restricted volume above the membrane. It is gratifying that the SpringSaLaD simulations produce these correct steady state surface densities even when, as in the last 2 rows of Table 1, mass action kinetics are not obeyed.

Table 2 provides results for three computational experiments in which structural features of the SpringSaLaD molecules are varied. These are all for dimerization reactions where the maximum distance between the membrane and the binding site is four times longer than in Table 1: *h* = 0.0205 µm (linker length of 20nm and binding site radius of 0.5nm). The 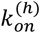 is therefore a factor of ~4 slower than that in Table 1 for the same 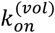 of 1µM^−1^s^−1^. The SpringSaLaD simulations were carried out with a slow membrane-anchor diffusion coefficient, *D*_*mem*_= 0.01µm^2^/s, to model a transmembrane protein domain. The first row can be directly compared to the fifth row of Table 1, where the only difference is the length of the linker. As would be expected from the increase in *h*, the membrane on rate constant is decreased; however, importantly, this constant is now closer to the diffusion rate and therefore deviations from mass action are significantly reduced. The second row in Table 2 shows results for a structure where additional spherical sites are introduced between the membrane anchor and the binding site to model the space occupied by a cytosolic protein sequence; this steric effect results in a small decrease in 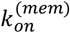. This decrease is reversed when flexibility is introduced by allowing the spherical sites to be pivot points in the third row of the Table; this is how disordered domains may be modeled in SpringSaLaD. Overall, for these idealized structures and simple dimerization, the fitted on-rate constants in Table 2 are relatively close to 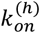.

### The appropriateness of mass action kinetics for reactions of binding sites tethered to a membrane.

Qualitatively, one can consider the overall rate coefficient, *k*_*on*_, to be affected by the intrinsic binding occurring once the sites are close enough to bind and characterized by the rate constant *k*_0_, and by diffusion-influenced reactant encounters characterized by their corresponding rate coefficient *k*_*D*_. In 3D, *k*_*D*_ is a time-independent constant (12), and so is *k*_*on*_, which in this case is computed as 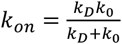 (13). As discussed in the Introduction and in the previous section in connection with the third panel of Figure 2, simple mass action kinetics are not always appropriate for bimolecular reactions within a surface because *k*_*D*_ becomes a function of time (8,12,13). Indeed, in the diffusion-limited regimes with *k*_*D*_ (*t*) ≪ *k*_0_, the *k*_*on*_ will be largely determined by *k*_*D*_ (*t*) and thus will depend on time, so a mass action rate law will be a poor fit to the kinetics. We now wish to analyze our tethered binder reactions for their conformance to mass action, using the criteria proposed by Yogurtcu and Johnson (10). Full mathematical details of our approach are found in Supporting Text II; we will provide a summary here.

For the dimerization of binding sites anchored to the membrane with a single simple tether, the intrinsic binding rate constant is 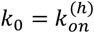. For cases of short tethers with lengths that are significantly less than the average initial distance between the anchors (exemplified by Table 1), the effective reaction radii are determined by tether lengths *h*, and the effective diffusion coefficient is determined by the diffusivity of anchors *D*_*mem*_. Also among the factors affecting possible deviations from simple mass action are the reactant densities (or the degrees of reaction completion), as discussed above in relation to the development of depletion zones. Thus, for a given combination of tether length, diffusion coefficient, effective 2D on-rate constant, initial surface density and desired degree of completion for a fit, we need to define a metric that describes how well *k*_*on*_ may be approximated as a constant (i.e. by 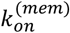).

As a first step, we needed to determine the general functional form of *k*_*D*_ (*t*), which is the source of the time dependence of *k*_*on*_. To do this, we solved the Smoluchowski model (12) numerically in 2D to produce the relation shown in Fig. 3. The results are plotted using dimensionless variables to allow us to determine *k*_*D*_ as a function of time for any combination of *D*′ (the sum of the *D*_*mem*_ for the two reactants) and *r*_*AB*_ (the sum of the radii of the two reactants). Full details of how Fig. 3 was calculated are found in Supporting Text II in relation to Fig. S7, where the calculations are also validated against short-time and long-time asymptotic analytical solutions.

With *k*_*D*_ (*t*) in hand, we can now ask how much it contributes to *k*_*on*_ for a desired level of completion, *p*, (e.g. 80% in Tables 1 and 2.); 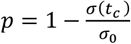, where *σ*(*t*) is the monomer surface density and *t*_*c*_ is the time to the desired level of completion, which can be read off the kinetic data for *σ* vs. *t*. (It is important to appreciate that *t*_*c*_ is smaller for higher initial surface density, all other parameters being equal; therefore, the range of *k*_*D*_ (*t*) that pertains is higher up the initial segment of the plot in Fig. 3, increasing the contribution of *k*_0_ to *k*_*on*_). In general, we can establish the relative importance of *k*_*D*_ (*t*) by defining the metric *δ* as

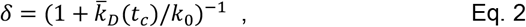

where 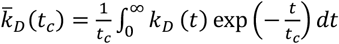 is, in effect, a weighted average of *k*_*D*_(*t*) over the time interval between *t* = 0 and *t* = *t*_*c*_. Thus, Eq 2 is indeed a measure of deviation from mass-action kinetics *k*_0_: if *δ* is close to 1, 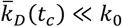, i.e. the reaction is diffusion-limited and *k*_*on*_ will not be constant; conversely, if *δ* is small, the kinetics will be dominated by *k*_0_ and will be well approximated by mass action. Eq. 2 is fully derived in Supporting Text II, where it is Eq. S11. Values of *δ* are included in Table 1 for all the irreversible reactions; these were calculated using 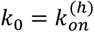 and *r*_*AB*_ = 2*h*, as discussed above (see also Supporting Text I). As illustrated in Figure 2, the parameters associated with row #5 of Table 1 produce kinetics in SpringSaLaD that are not approximated well by the mass-action rate law; the corresponding value of *δ*, 0.78 is indeed close to 1. We judge a value of *δ* < 0.3 to allow for the application of 2D mass action as a reasonable approximate rate law, but individual judgment should be made on how critically the overall system behavior might depend on conformity to mass action kinetics.

In Supporting Text II, we also show that a weighted average of the overall on-rate constant, 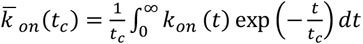 can be computed as

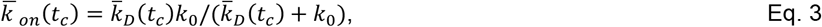

which is structurally similar to the equation for *k*_*on*_ in 3D. Using Eq. 3, we obtained a gratifyingly good match for 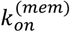, derived by fitting the SpringSaLaD simulations.

In the case of long tethers, 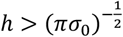, exemplified by Table 2, geometric constraints imposed by tethers and slow anchor diffusion are less limiting in two respects. First, multiple binding sites may now encounter each other under the hemisphere of radius *h*. As a result, the tether length no longer represents the effective reaction radius, which should now be determined by the assigned radii of the binding sites 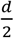. Second, the main mixing mechanism is now volumetric diffusion. Because the area of the hemisphere is twice the area of its projection on the membrane, and both are covered by diffusion within the same time, the corresponding effective 2D diffusivity is 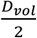. Thus, to compute δδ from Eq. 2 for the case of long tethers, we should compute *k*_*D*_ (*t*) from the relation of Fig. 3 with *r*_*AA*_ = *d* and *D*^′^ = *D*_*vol*_. The resultant *δ* for the first row of Table 2 is 0.03, indicating that mass action is a good approximation. The other rows of Table 2 also are clearly well fitted by mass action (see the fits in the spreadsheets in Supporting Information), but we did not calculate *δ* because 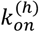 is not as well defined when the structures become more complex than a single tethered sphere. This is because both steric effects and excluded volume effects will impact binding kinetics in these cases.

In general, metrics like *δ* will not be easy or possible to calculate for complex structures based on real molecules. Thus, in general the best way to assess the applicability of mass action as a reasonable approximation for membrane-tethered reactions is to judge the fit of the SpringSaLaD simulation to the integrated mass-action rate law. As can be seen by comparing the plots in Fig. 2 and the corresponding entries in Table 1, it is clear that the fits for rows 1 and 6 are good, while the fit for row 5 is poor, consistent with both the entries for *δ* and the relative squared deviation of the fit. But even a poor fit to mass action may be an adequate approximation for some situations, especially where the uncertainty in other parameters of a large signaling system can be large. Additionally, for reversible binding, the steady state distribution is not dependent on the accuracy of mass action kinetics. We now illustrate our method by applying it to a real signaling module that has been widely investigated because of its importance in cancer biology.

### Application of the method to interaction of receptor-bound SOS with Ras.

Until now, we have employed idealized molecular structures to validate our method and to learn more about biophysical principles that control the on-rate constants of binding sites tethered to membranes. We now illustrate the application of this approach to a biologically relevant example, namely the interaction of the G-protein exchange factor (GEF) SOS with the lipid anchored small G-protein Ras (22).

SOS has two binding sites for Ras: an allosteric site and a catalytic site. When a Ras molecule binds to the allosteric site it increases the GEF activity of the catalytic site (36). Additionally, before SOS binds to Ras it is first recruited to an active receptor tyrosine kinase (RTK) through an adapter protein; the adaptor binds to a proline rich motif (PRM) on SOS via a SH3 domain and to a phosphorylated tyrosine via a SH2 domain. One such RTK is the Epidermal Growth Factor Receptor (EGFR) and one such adaptor protein is Grb2 (37). Once SOS is bound to Grb2, it becomes membrane-tethered and its interaction with Ras is facilitated (36). However the complex mechanistic details of this system are still emerging (38).

We asked the limited question of how binding of Ras with the receptor-associated SOS catalytic site might depend on whether SOS is prebound to Ras at the allosteric site. Ras is a lipid-anchored protein, so we reasoned that binding of Ras to the allosteric site of SOS would bring the SOS catalytic site closer to the membrane to enhance binding to a second Ras and subsequent exchange of GDP for GTP. Just how large an effect this is, may be estimated by the procedure developed above, with the results shown in Fig. 4.

We developed molecular models with the aid of the mol2sphere (25) utility within SpringSaLaD and were guided by AlphaFold 2 atomic structure predictions (23,24); all the site diameters and linker lengths are available in the SpringSaLaD input file included in the Supporting Information; snapshots of the structure are shown in Fig. 4. The top of Fig. 4 displays results for binding of SOS-Grb2-EGFR to Ras at the SOS catalytic site; the bottom shows results for the same reaction, except SOS-Grb2-EGFR had been first bound to a Ras molecule at the SOS allosteric site. The input rates shown at the top of Fig. 4 are based on experimentally measured data (22) and are applied to both of the reactions. Special considerations were applied to ensure accuracy of the on rate, which was parameterized via an initial cytosolic reaction where the output rate constant was matched to the experimentally measured rate data (22). This important step allows us to calibrate the coarse-grained protein structure used in our model to real protein structures, which inherently impact binding *in-vivo* and are therefore reflected in experimentally measured reaction rates. The desired output rate corresponding to an experimentally observed rate constant of 0.27 µM^−1^s^−1^ (22), was achieved with an input rate constant of 0.35 µM^−1^s^−1^, which was then used in subsequent simulations of membrane bound interactions in Fig. 4. In these models, the EGFR membrane anchor site is assigned a diffusion coefficient of 0.01 µm^2^/s to represent a large transmembrane protein, while the Ras membrane anchor is assigned a diffusion coefficient of 1.0 µm^2^/s to represent a lipid anchor; all the sites that are dangling in the cytosol volume are given *D*_*vol*_ of 1.0 µm^2^/s, but since the binding reaction is not close to diffusion-limited, the precise values are not critical. Consistent with these being reaction-limited on-rates, the SpringSaLaD simulation outputs (averages of 100 runs) are very well fitted to reversible mass action kinetic law, as shown in the plots on the right of Fig. 4 (relative squared deviations are, respectively, 4.1 × 10^−4^ and 2.0 × 10^−4^). Importantly, the 2D on-rate constants (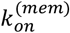) derived from these fits are, respectively, 5.0 × 10^−3^ µm^2^molecules^−1^s^−1^ and 1.9 × 10^−2^ µm^2^molecules^−1^s^−1^. Likewise, the affinity of the catalytic site is increased by allosteric site pre-association: *K*_*d*_ = 800 molecules/µm^2^ for the top of Fig. 3 and 210 molecules/µm^2^ for the bottom pre-association case. Thus, SOS allosteric site association with Ras is estimated to accelerate its catalytic site binding and affinity by a factor of ~4 – even when SOS is already confined to the membrane through Grb2-mediated association with EGFR.

### A simple ODE model of receptor-mediated signaling

To illustrate the application of our approach, we built a small signaling model based on the values of 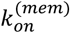 determined in the previous section. The ODE model was built in VCell (29,30) and total concentrations of the various species were from measurements on HeLa cells (26,27). In addition to the two reactions from Fig. 4, we also modeled recruitment of Grb_SOS from the cytosol to the membrane by phosphorylated EGFR (EGFRPTyr) and direct stepwise binding of two Ras molecules to Grb_SOS; the model assumes that almost all the cytosolic SOS is prebound to Grb2, which is present in excess (39). The VCell reaction diagram in Fig. 5A displays the connectivity of the network and Table 3 provides the corresponding mass action rate constants.

We used the data for EGFR measured in HeLa cells (26) to derive an initial surface density for EGFPtyr of 840 molecules/µm^2^, which we took to be the fully activated receptor (we did not attempt to account for kinetics of inactivation, which would lower this number, or for the multiple phosphorylation sites available on the cytoplasmic domain of EGFR, which would raise this number). Fig. 5B shows the approach to steady state for this “Full activation” condition. It shows that free EGFPtyr and free Ras are only partially depleted at steady state, while ~90% Grb_SOS is removed from the cytosol to become membrane associated. Figure 5C provides a more detailed look at the association of Grb_SOS with Ras. We chose three levels of initial EGFRPTyr, 0, 200 and 840 molecules/µm^2^, to model, respectively, basal unstimulated cells, partial activation of EGFR and full activation. The species composed of doubly bound Ras, Grb_SOS_Ras_Ras and EGFR_Grb_SOS_Ras_Ras (GSRR and EGSRR in Fig. 5), are always a factor of ~3.5 higher than the corresponding species with one Ras. This is important because the binding two Ras molecules is required for SOS to optimally express its GEF activity (40); the second binding event is, of course, significantly enhanced by the proximity effect quantified in Fig. 4; this enhancement at the membrane has been reported based on experiments for the binding of Ras to SOS alone (41). Another conclusion from Fig. 4C is that the overall levels of Ras doubly bound to SOS are significantly increased by EGFR activation; this is consistent with the high affinity of the SH2 domains on Grb2 for phosphotyrosine (42), creating a higher local concentration of SOS near the membrane surface. Indeed, under the “Basal” condition without participation of EGFRPtyr, 80% of the Grb_SOS remains unbound in the cytosol (data not shown but available in the public VCell Model “SOS_Recruitment_Ras_Binding”).

## Discussion

The kinetics of reactions at membranes have long fascinated biophysicists (1,5,7,8,10,11,17,19,32). These studies have produced theoretical insights to illuminate how surface-associated reactions have distinct properties compared with reactions occurring in 3D solution. Which of these special properties are most pertinent to any given membrane-bound molecular interaction is difficult to ascertain *a priori*. Furthermore, experiments to measure bimolecular kinetics on membrane surfaces are complex (4), so often only on-rate constants measured in 3D are accessible. Fundamentally, however, the kinetics of key membrane-associated reactions depend on the 2D surface densities and 2D rate constants, not on the bulk cellular concentrations and 3D rate constants. Indeed, because surface to volume ratios of different cell types vary tremendously, volumetric rate constants cannot be readily used to model and simulate cell signaling systems. To address these theoretical and practical problems, we describe a procedure (Fig. 1) using experimentally accessible volumetric on-rate constants, 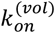 within the SpringSaLaD simulation software to estimate the 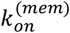, the 2-dimensional rate constant for a membrane-confined bimolecular reaction.

To validate the method, we applied it to the dimerization of a single binding site tethered to a surface through a 5nm stiff linker, where the membrane anchor acts as a pivot (Table 1). For the situation where the reaction is rate limiting, this system can be solved theoretically (see Supporting Text I); gratifyingly, 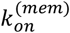 determined by our method is well reproduced by this solution. Interestingly, for these cases, 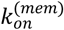 is well approximated by 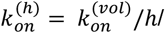602.2 (µm^2^molecules^−1^s-^1^), where *h* is the distance of the binding site from the membrane anchor (in µm) and 602.2 is a units conversion factor. The parameter *h* has also been referred to as the “confinement length” (17), defining a thin volume above the membrane that concentrates the binding sites and directly producing the relationship between 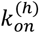 and 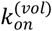.

While mass action kinetics are generally applicable for both encounter-limited and reaction-limited kinetics in 3D solution (but see ((12)15)), it has long been appreciated that the situation may be more complex for 2D kinetics (5,8,10,11,32,43). This is demonstrated by the results in Table 1 for situations where the diffusion coefficient of the anchor is slow, but the volumetric on-rate constant is fast. For these cases, different estimates of 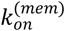 are obtained at different initial surface density – clearly incompatible with the mass action kinetics. Indeed, the third panel of Figure 2 shows that the SpringSaLaD kinetic data is not well fit by a 2D mass action rate law. A video of one of these trajectories (Movie 1) nicely illustrates how the initial rate is fast, while the binding sites are within “reach” (19), but falls off as binding sites are left orphaned outside the reach of the remaining slowly diffusing monomers. Because of this, a mass action rate law applies better to high initial concentrations and to shorter time durations, before depletion zones develop. To address the applicability of mass action theoretically, we solved the Smoluchowski model (12) in 2D using VCell (Supporting Text II) to generate the relation plotted in Fig. 3 for the time-dependence of the diffusion limited reaction coefficient. This allowed us to implement a pipeline for computing a weighted average of the on-rate coefficient over the time to a particular level of reaction completion, for any intrinsic on-rate, *k*_0_, membrane diffusion coefficient, initial surface density, and desired level of completion for a fit. We used this to see how well the idealized scenarios in the first 8 rows of Table 1 and the first row of Table 2 could be fitted by simple mass action (i.e. where the rate coefficients can be reasonably well approximated as independent of time). We were able to derive a metric, *δ*, that measures how significantly the overall rate is influenced by diffusion. This is similar to the detailed analysis of Yogurtcu and Johnson (10) for 2D reactions, except their analysis focused on pure 2D (i.e. *h* = 0). Our results allow us to generalize that mass action applies as long as 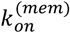 is close to 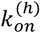. Importantly, there was a clear relationship between how well the SpringSaLaD simulation results can be approximated with a mass action rate law, as measured by the relative squared deviations and *δ* in Table 1. Thus, our analysis provides an approach to determine whether 2D binding might be well approximated by mass action kinetics, and if so, to estimate the 2D mass action on-rate constant.

To explore how other molecular structural features might affect dimerization of the monomers tethered to the membrane, we looked at three additional idealized systems in Table 2. In all these, *h* was 20.5nm (as opposed to 5.5nm in Table 1). As expected, the longer confinement length decreased the estimated 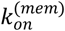 by a factor of ~4. The insertion of steric sites between the anchor and the binding site or allowing for flexibility of the linker region have minor effects on 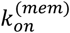, which is relatively well approximated by 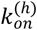. Importantly, the lower value of 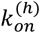 for this system was closer to *D*_*mem*_, resulting in a better fit by the mass action kinetics than for the similar case with the 5nm linker (fifth row of Table 1).

For membrane binding of real biological molecules, the average location of binding sites relative to the membrane surface (i.e. *h*) will generally be not well defined, since it is influenced by the steric effects and the variable flexibility of neighboring protein domains. Also, the two binding sites may be parts of very different structures with different distances from the membrane surface. In situations like this, our approach has the potential to provide good estimates of rate constants that can be applied to larger cell signaling systems. Indeed, there may be direct insights that can be realized just by considering the structural details of the interacting membrane molecules. We have illustrated this in relation to adaptor-mediated protein kinase receptor signaling mechanisms, specifically for the interaction of the GEF SOS with its effector Ras (Fig. 4); the fit of the SpringSaLaD simulations to a mass action rate law is excellent in both scenarios of Fig. 4, indicating that mass action is applicable. It has been shown that direct catalysis by the SOS catalytic domain of Ras conversion from the GDP to the GTP states is relatively slow. However, binding of Ras to SOS at a site that is not catalytic (termed the “allosteric” site on SOS) significantly accelerates the catalytic activity(40,41), where the catalysis becomes processive (22,36,38). The results in Fig. 4 suggest that at least part of this acceleration may be due to the close proximity of the SOS catalytic site to the membrane once it is bound to Ras at its allosteric site. Even though SOS is already localized to the membrane by initially binding to EGFR via Grb2 in our computational experiment, pre-binding of the SOS allosteric site to Ras brings it even closer to the membrane. Of course, there could be additional effects such as a direct allosteric enhancement through a conformational change or release of self-inhibition (22,36,38), but here we focus on the significance of constricting the binding zone through membrane tethers of varying length and flexibility.

To illustrate the application of our approach to cell signaling, we constructed a simple model of EGFR signaling to Ras via SOS in HeLa cells (Fig. 5). The results show that the proximity effect established in Fig. 4 for the binding of SOS to the second Ras molecule results in the dominance of doubly bound Ras species. This no doubt contributes to the efficiency of the GEF activity associated with SOS. Our simple model also demonstrates the effectiveness of active EGFR in boosting the recruitment of Grb_SOS to the membrane where it presents an effectively high local concentration to Ras. The model, which is available in the VCell database, can be further explored for the effect of varying Ras or SOS concentrations or the behavior of different cell types with different levels of the key signaling models; it could also be used as a starting point for more elaborate models that incorporate more detailed mechanisms and additional downstream signaling events.

Thus, our approach toward estimating membrane on-rates will aid in the parametrization of ODE and PDE models that could help elucidate the full kinetic and mechanistic details of cell signaling systems. In ODE models, diffusion is assumed to be fast on the timescale of reaction rates; however, our approach is necessary to capture the special local membrane effects that allow conversion of a bulk measurement of the on-rate to 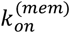. In cell-scale PDE (or spatial stochastic) models, where diffusion is considered explicitly, the membrane molecules are treated as points and the distances are very large on the scale of molecular dimensions. So the most appropriate diffusion coefficient would be that of the membrane anchor. These are typically measured experimentally using fluorescence recovery after photobleaching (FRAP) or fluorescence correlation spectroscopy (FCS). It is also possible to estimate membrane diffusion coefficients based on the size of the membrane embedded region and the known viscosity of biological membranes.

## Supporting information

Derivations for limiting cases related to Table 1 (Supporting Text I) and Derivations related to Applicability of mass-action kinetics (Section II)

Movie of an example trajectory for the model of Table 1 Row 5 (and also Fig. 2, 3rd panel).

## Author contributions

KJP created the computational models, performed the simulations, analyzed the data and wrote the paper; BMS performed the calculations in the Supporting Text I and II and wrote the paper; LML conceived the research, analyzed the data and wrote the paper.

## Acknowledgments

This work was supported by NIH grants R24 GM137787 and R01 GM132859. We are pleased to acknowledge the advice of Aniruddha Chattaraj with some of the data analysis.

## List of Supporting Data and Information

Derivations for limiting cases related to Table 1 (Supporting Text I) and Derivations related to Applicability of mass-action kinetics to dimerization of molecules with binding sites tethered to the membrane. Filename: Supporting Text_revised.docx

SpringSaLaD input files for the models in Tables 1 and 2 and Figure 4. Filenames: Table 1 and 2 SIMS.zip; EGFR_Grb_SOS binding to Ras.txt; EGFR_Grb_SOS_prebound at allo binding to Ras.txt Spreadsheets related to Tables 1 and 2 and Figure 4 containing the SpringSaLaD simulation results and the fits to determine 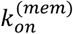. Filenames: Table 1 5nm_stiff_fits.xlsx; Table2 20nmT_fits.xlsx; Data for Figure 4.xlsx

Movie of an example trajectory for the model of Table 1 Row 5 (and also Fig. 2, 3^rd^ panel). Filename: Movie kon=1 Danchor=membrane protein.mp4

