## Supplementary material for "Mesoscale simulations of membrane-tethered reactions to parameterize cell-scale models of signaling": Derivations for limiting cases related to Table 1 (Supporting Text I) and Derivations related to Applicability of mass-action kinetics (Section II)

**I. Derivation of an equivalent 2D dimerization rate constant for tethered binders in the limit of** $\boldsymbol{D}_{\boldsymbol{mem}}\boldsymbol{\ll}\boldsymbol{D}_{\boldsymbol{vol}}$ **.**

Here we derive $k_{on}^{\left( mem \right)}$ analytically for conditions pertaining to the first row in Table 1 of the main text. Our approach is conceptually similar to the one previously used in modeling the binding of bivalent haptens to an antibody (Dembo and Goldstein, 1978).

As in SpringSaLad (Michalski and Loew, 2016), we model a binder as a ball of diameter $d$. A link of length $L$ tethers the ball’s center to a membrane-bound anchor. Because the anchor diffuses in the membrane much slower than the binder in the cytosol, the link restricts movements of the binder, so that at any given time, the binder is generally confined to a hemisphere, centered at a current position of the anchor, with the radius $R=L+\frac{d}{2}$ (in the main text, this parameter is termed $h$). The simulation results, which we seek to understand, were obtained for $d<\frac{1}{5}L$, so that the excluded volume effects due to the binder’s finite size can be ignored. We disregard possible effects on $k_{on}^{\left( mem \right)}$of the collisions or entanglement of the tethers, given that the links in SpringSaLad are not affected by each other.

Importantly, we assume that during dimerization, the monomers remain uniformly distributed. A formal analysis of the conditions underlying this assumption is provided in part II below.

With the assumptions outlined above, the effective 2D rate constant $k_{on}^{\left( mem \right)}$ for the dimerization of monomers tethered to the membrane can be obtained analytically in terms of the dimerization rate constant $k_{on}^{\left( vol \right)}$ of free binders.

Two binders can collide if the distance between their anchors does not exceed $2R$. As mentioned earlier, the anchors can be viewed as “well-mixed” with some surface density $\sigma$. Without loss of generality, we may assume that the anchor of one of the monomers is immobile (in what follows, we will call such a monomer ‘a given monomer’). The number of binding partners of this monomer is then

$n=$ $4\pi R^{2}\sigma$.	Eq (S1)

Our goal is to derive the rate of dimerization of a given monomer.

We first consider cases where the links between a binder and its anchor are stiff. In such cases, given that $D_{mem}\ll D_{vol}$, the binder may be thought of as uniformly distributed at any time within a hemispherical shell centered at a current position of its anchor. The shell has the outer radius $R$, the inner radius $r=L-\frac{d}{2}$, and the volume $v_{shell}=\frac{2\pi R^{3}}{3}\left( 1-\left( \frac{r}{R} \right)^{3} \right)$ , so the binder ‘concentration’ (more precisely, the binder probability density function) is $\frac{1}{v_{shell}}=[\frac{2\pi R^{3}}{3}{\left( 1-\left( \frac{r}{R} \right)^{3} \right)]}^{-1}$.

The dimerization of a given monomer with a binding partner requires that their binders collide, which occurs at the intersection of their shells (the shared space in Figure S1). The collision probability for the binder of the given monomer, distributed within the shell shaded in Figure S1, $p_{coll}=\frac{v_{shared}}{v_{shell}}$ , where $v_{shared}$ is the volume of the shared space. As $v_{shared}$ depends on distance $l$ separating the anchors, so does the collision probability. Because the anchors are well-mixed, the mobile anchor of the binding partner (the center of the unshaded shell in Figure S1) is uniformly distributed within the circle of radius $2R$ with the center at the fixed anchor. The average collision probability is found by integrating of $p_{coll}\left( l \right)=\frac{v_{shared}\left( l \right)}{v_{shell}}$ over this circle:

**Figure S1. Cross-section of overlapping hemispherical shells.**

The shells cover binder positions of two monomers with anchors separated by distance $l$.

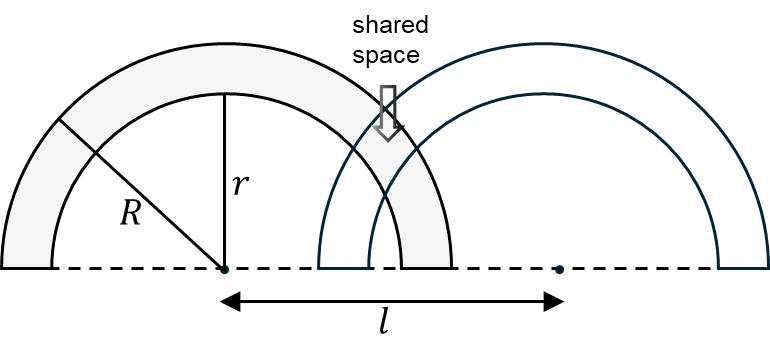

$\bar{p}_{coll}=\frac{1}{\pi\left( 2R \right)^{2}}\int_{0}^{2\pi} d\varphi\int_{0}^{2R} p_{coll}\left( l \right)ldl=\frac{2}{\left( 2R \right)^{2}}\int_{0}^{2R} p_{coll}\left( l \right)ldl$. Eq (S2)

Then the rate of dimerization of the given monomer with a single binding partner is the product of the rate constant of dimerization of free binders $k_{on}^{\left( vol \right)}$ and the ‘concentration’ of the single binder $\frac{1}{v_{shell}}$ modified by the average collision probability: $\bar{p}_{coll}\frac{k_{on}^{\left( vol \right)}}{v_{shell}}$.

We now recall that the given monomer has $n$ binding partners defined by Eq (S1). For anchor densities that are not too high, so that the events of three-binder collisions can be ignored, the rate of dimerization of a given monomer with any of the available binding partners is simply $n\bar{p}_{coll}\frac{k_{on}^{\left( vol \right)}}{v_{shell}} =\left( \frac{4\pi R^{2}\bar{p}_{coll}k_{on}^{\left( vol \right)}}{v_{shell}} \right)\sigma$, where the expression in the parenthesis is the sought formula for the equivalent $k_{on}^{\left( mem \right)}$:

$k_{on}^{\left( mem \right)}=$ $\frac{4\pi R^{2}\bar{p}_{coll}k_{on}^{\left( vol \right)}}{v_{shell}}= \frac{4\pi R^{2}}{v_{shell}^{2}} k_{on}^{\left( vol \right)} \int_{0}^{1} v_{shared}\left( \rho\right)2\rho d\rho$, Eq (S3)

where we introduced the dimensionless variable $\rho=\frac{l}{2R}$ .

We now turn to calculations of $v_{shared}\left( \rho\right)$. Introducing for brevity the notation $a=\frac{r}{R}$, we observe that for any $a,\rho\leq1$, the space shared by two intersecting spherical shells is a combination of spherical caps. It is therefore convenient to introduce a function,

$f\left( x \right)=1-\frac{3}{2}x +\frac{1}{2}x^{3}$,	Eq (S4)

which for $x=\cos\theta$ yields the volume fraction of a hemisphere that is occupied by a spherical cap with polar angle $\theta$ (Figure S2).

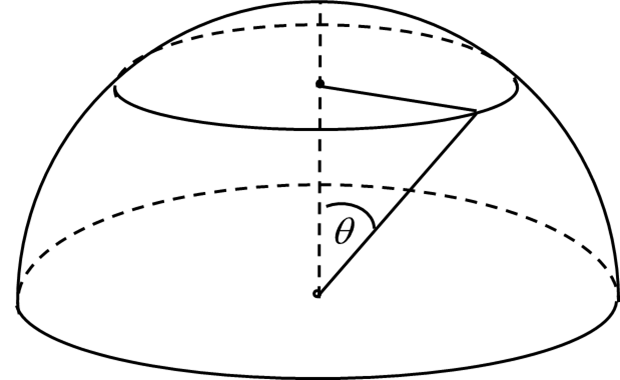

**Figure S2. A spherical cap defined by a polar angle** $\boldsymbol{\theta}$ **.**

The cap is shown as a part of the corresponding hemisphere.

The structure of $v_{shared}\left( \rho\right)$ depends on the value of $a$. For $a\geq\frac{1}{3}$, the derivation based on stereometric considerations yields the following dependence:

$\frac{v{}_{shared}\left( \rho\right)}{\left( 2\pi R^{3}/3 \right)}=\left\{ \begin{aligned} \begin{matrix} f\left( \rho\right)-a^{3}f\left( -\frac{\rho}{a} \right),\mathrm{for} \rho\in\left[ 0, \frac{1}{2}\left( 1-a \right) \right] \\ f\left( \rho\right)+a^{3}f\left( \frac{\rho}{a} \right)-f\left( \frac{1-a^{2}}{4\rho}+\rho\right)-a^{3}f\left( \frac{a^{2}-1}{4\rho a}+\frac{\rho}{a} \right),\mathrm{for} \rho\in\left[ \frac{1}{2}\left( 1-a \right), a \right] \\ f\left( \rho\right)-f\left( \frac{1-a^{2}}{4\rho}+\rho\right)-a^{3}f\left( \frac{a^{2}-1}{4\rho a}+\frac{\rho}{a} \right),\mathrm{for} \rho\in\left[ a, \frac{1}{2}\left( 1+a \right) \right] \end{matrix} \\ f\left( \rho\right),\mathrm{for} \rho\in\left[ \frac{1}{2}\left( 1+a \right), 1 \right] \end{aligned} \right.$ . Eq (S5)

Eq (S5) applies to the stiff-tether cases discussed in the main text. Indeed, for $L=5 \mathrm{nm}$ and $d=1 \mathrm{nm}$,the outer and inner shell radii are $R=5.5 \mathrm{nm}$, $r=4.5 \mathrm{nm}$, resulting in $a=\frac{r}{R}=\frac{9}{11}>\frac{1}{3}$, and

the monomers with larger $L$ and same $d$ are characterized by even larger $a$. Figure S3

illustrates $\frac{v_{shared}}{\frac{2\pi R^{3}}{3}}$ as a function of $\rho$ for $a=\frac{9}{11}$.

**Figure S3. Volume of intersection of two identical shells as a function of distance between their centers.**

The volume of the shared space, normalized to the volume of the hemisphere, is shown as a function of the normalized distance $\rho=\frac{l}{2R}$; $L=5 \mathrm{nm}, d=1 \mathrm{nm}$.

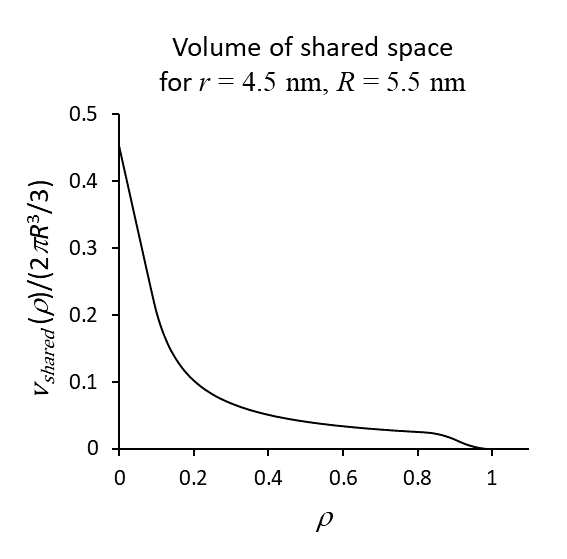

The integral in Eq(S3) is readily evaluated,

$\int_{0}^{1} v_{shared}\left( \rho\right)2\rho d\rho=\frac{2\pi R^{3}}{3}\{\frac{1}{5}\left( 1+a^{5} \right)+\frac{a}{8}\left( 1-a^{2} \right)^{2}-\frac{1}{80}(\left( 1+a \right)^{5}-\left( 1-a \right)^{5})\}$. Eq (S6)

Substituting Eq (S6) into Eq (S3) and noting that $v_{shell}=\frac{2\pi R^{3}}{3}\left( 1-\left( \frac{r}{R} \right)^{3} \right)=\frac{2\pi R^{3}}{3}\left( 1-a^{3} \right)$, we find

$k_{on}^{\left( mem \right)}=\phi\left( a \right)\frac{k_{on}^{\left( vol \right)}}{R},$	Eq (S7a)

where	$\begin{matrix} \\ \phi\left( a \right)=\frac{\frac{6}{5}\left( 1+a^{5} \right)+\frac{3a}{4}\left( 1-a^{2} \right)^{2}-\frac{3}{40}\left( \left( 1+a \right)^{5}-\left( 1-a \right)^{5} \right)}{\left( 1-a^{3} \right)^{2}} \end{matrix}$ . Eq (S7b)

In the terms of the main text, $\frac{k_{on}^{\left( vol \right)}}{R}=\frac{k_{on}^{\left( vol \right)}}{h}=k_{on}^{\left( h \right)}$, and from Eqs (S7a), $\phi\left( a \right)=\frac{k_{on}^{\left( mem \right)}}{k_{on}^{\left( h \right)}}$. For stiff tethers with length $L=5 \mathrm{nm}$ and a binder’s diameter $d=1 \mathrm{nm}$, $a=\frac{9}{11}$ and $\frac{k_{on}^{\left( mem \right)}}{k_{on}^{\left( h \right)}}=1.06$. This is close to $\frac{k_{on}^{\left( mem \right)}}{k_{on}^{\left( h \right)}}=0.967$ from the first row of Table 1 (main text). Thus, the analytical solution validates the simulation results obtained with SpringSaLaD.

Note that the ratio $\frac{k_{on}^{\left( mem \right)}}{k_{on}^{\left( h \right)}}$ are not particularly sensitive to tether lengths $L$ in the limit $D_{mem}\ll D_{vol}$ (Figure S4A), and it is nearly the same as in the case of $D_{mem}=D_{vol}$ (second row of Table 1). This appears counterintuitive, given that for $D_{mem}\ll D_{vol}$, the tethering causes the collision probability to decrease with $L$, whereas for equal diffusivities, the binders are effectively unconstrained by their anchors in their movements within the layer of height $L+\frac{d}{2}$, adjacent to the membrane.

**Figure S4. Ratio** $\boldsymbol{k}_{\boldsymbol{on}}^{\boldsymbol{(mem)}}\boldsymbol{/}\boldsymbol{k}_{\boldsymbol{on}}^{\boldsymbol{(h)}}$ **has low sensitivity to tether length** $\boldsymbol{L}$ **.**

(A) The ratio $k_{on}^{(mem)}/k_{on}^{(h)}$ as a function of the ratio of binder diameter $d$ and tether length $L$; $d=1 \mathrm{nm}$. (B) Interplay of main determinants of $k_{on}^{(mem)}/k_{on}^{(h)}$ as functions of $d/L$. (C) The log-log plot of collision probability as a function of binder ‘concentration’ in the shell, indicating that they are nearly reciprocal; this explains the low sensitivity of $k_{on}^{(mem)}/k_{on}^{(h)}$ to $L$.

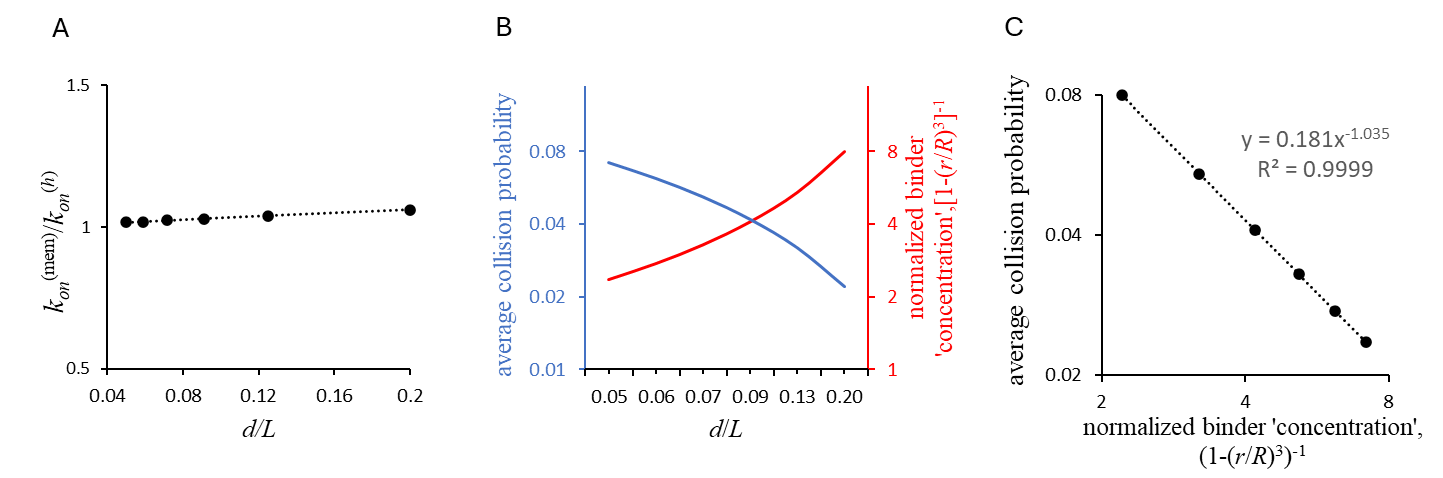

The low sensitivity of the coefficient of proportionality between $k_{on}^{\left( mem \right)}$ and $k_{on}^{\left( h \right)}$ to $L$ in the limit $D_{mem}\ll D_{vol}$ is explained by the interplay of the collision probability and the normalized binder ‘concentration’ within the shell, another determinant of $\frac{k_{on}^{\left( mem \right)}}{k_{on}^{\left( h \right)}}$ (Figure S4B). Indeed, these factors

vary with $\frac{d}{L}$ in a nearly reciprocal manner (Figure S4C).

That the value of $\frac{k_{on}^{\left( mem \right)}}{k_{on}^{\left( h \right)}}$ in the limit $D_{mem}\ll D_{vol}$ is close to that in the case of equal diffusivities is largely due to the assumption of well-mixed anchors: the averaging of $p_{coll}\left( l \right)=\frac{v_{shared}\left( l \right)}{v_{shell}}$ is done over the uniform anchor distribution (Eq (S3)), and the number of binding partners (Eq (S1)) assumes that the anchors are well-mixed.

We now turn to cases of flexible tethers, again assuming the diameter of a binder $d$ to be significantly less than the tether length $L$. If a tether is totally flexible, then the entire hemisphere, centered at the anchor, is accessible to the binder, so we have a case of $r=0$ and, consequently, $a=0.$ For completeness, we will consider a ‘semiflexible’ tether model, where a portion of the tether adjacent to the anchor is stiff, whereas the remaining segment is flexible, so that the binder again is found with equal probability in a hemispherical shell with the inner radius $r$, though in this case $\left( R-r \right)\neq d$ .

We recall that Eq (S5) holds only for $a\geq\frac{1}{3}$. For $a<\frac{1}{3}$, the stereometry yields the following equation for $v_{shared}/(\frac{2\pi R^{3}}{3})$,

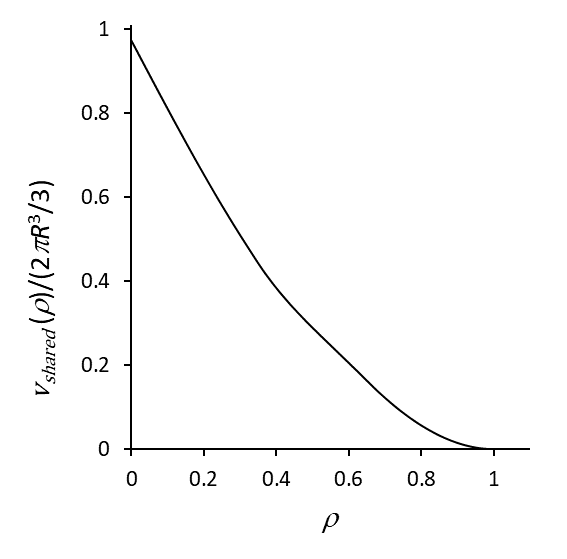

**Figure S5. Volume shared by two identical shells as a function of distance between their centers.**

The volume of the shared space, normalized by volume of the hemisphere, is shown as a function of $\rho=\frac{l}{2R}$; $\frac{r}{R}=0.3.$

$\frac{v{}_{shared}\left( \rho\right)}{\left( 2\pi R^{3}/3 \right)}=\left\{ \begin{aligned} \begin{matrix} f\left( \rho\right)-a^{3}f\left( -\frac{\rho}{a} \right),\mathrm{for} \rho\in\left[ 0, a \right] \\ f\left( \rho\right)-2a^{3}, \mathrm{for} \rho\in\left[ a, \frac{1}{2}\left( 1-a \right) \right] \\ f\left( \rho\right)-f\left( \frac{1-a^{2}}{4\rho}+\rho\right)-a^{3}f\left( \frac{a^{2}-1}{4\rho a}+\frac{\rho}{a} \right),\mathrm{for} \rho\in\left[ \frac{1}{2}\left( 1-a \right), \frac{1}{2}\left( 1+a \right) \right] \end{matrix} \\ f\left( \rho\right),\mathrm{for} \rho\in\left[ \frac{1}{2}\left( 1+a \right), 1 \right] \end{aligned} \right.$ . Eq (S8)

Figure S5 illustrates $v_{shared}/(\frac{2\pi R^{3}}{3})$ as a function of $\rho$ for $a=0.3$.

Interestingly, while Eq (S8) differs from Eq (S5), its integral is the same as in Eq (S6), so Eqs (S7) hold for all $a\in\left[ 0, 1 \right].$ Thus, for the case of ‘semiflexible’ tethers, the ratio $k_{on}^{\left( mem \right)}/(\frac{k_{on}^{\left( vol \right)}}{R})$ as a function of $a=\frac{r}{R}$ is described by Eq (S7b); the graph of the ratio as a function of a flexible fraction of the tether is shown in Figure S6.

For the fully flexible tether, it follows from Eq (S7b) that $\phi\left( 0 \right)=1.2$, therefore $k_{on}^{\left( mem \right)}=\phi\left( 0 \right)\frac{k_{on}^{\left( vol \right)}}{R}=1.2k_{on}^{\left( h \right)}.$

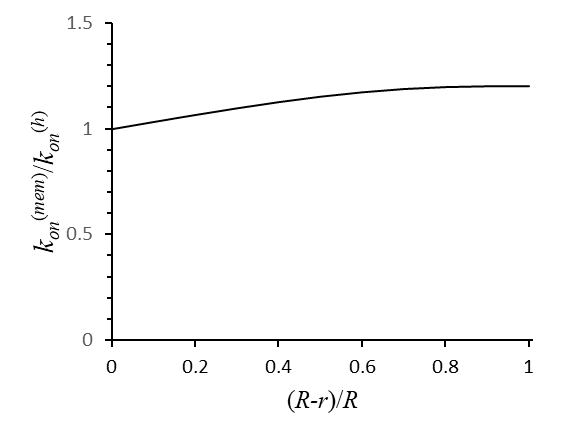

**Figure S6. Ratio** $\boldsymbol{k}_{\boldsymbol{on}}^{\boldsymbol{(mem)}}\boldsymbol{/}\boldsymbol{k}_{\boldsymbol{on}}^{\boldsymbol{(h)}}$ **for the case of monomers with (semi)flexible tethers.**

The ratio is shown as a function of a flexible fraction of the tether.

**II.** **Applicability of mass-action kinetics to dimerization of molecules with binding sites tethered to the membrane.**

Here we discuss the conditions under which the rate of dimerization of monomers tethered to the membrane can be accurately approximated with a mass action rate law, $k\sigma_{A}\sigma_{B}$, where $\sigma_{A}\left( t \right)$ and $\sigma_{B}\left( t \right)$ are the time-dependent two-dimensional (2D) densities of the binding partners $A$ and $B$, and $k$ is the rate constant, i.e. it is independent of time. In the case of homodimerization, the corresponding rate equation, $\partial_{t}\sigma=-2k\sigma^{2}$, has the exact solution,

$\sigma\left( t \right)=\frac{\sigma_{0}}{1+2k\sigma_{0}t}$,	Eq (S9)

where $\sigma_{0}$is the initial monomer concentration, $\sigma_{0}=\sigma\left( 0 \right)$.

We first consider the case of short tethers, such that their lengths are shorter than half the average initial distance between the anchors, $h=L+\frac{d}{2} <\left( \pi\sigma_{0} \right)^{-\frac{1}{2}}$. This case is exemplified by Table 1 of the main text. Note that the limit $L\to0$, that describes the binding of molecules embedded in the membrane, was studied by Yogurtcu and Johnson (Yogurtcu and Johnson, 2015).

As we showed in part I of this Supplement, the conditions under which the binding partners remain well-mixed result in mass-action kinetics. In the case of short tethers, the mechanism of monomer mixing is the diffusion of their anchors. Thus, it is qualitatively clear that the mass-action rate law would apply to reaction-limited binding, which is characterized by high anchor diffusivity and slow association of the binding sites upon collision. Under these conditions, the mass-action rate constant $k$ essentially coincides with the rate constant of intrinsic binding, $k_{0}$, (which corresponds to $k_{on}^{\left( h \right)}$ for the case of the tethered binding sites in part I). Such clear reaction limited binding is exemplified by the second row of Table 1 of the main text.

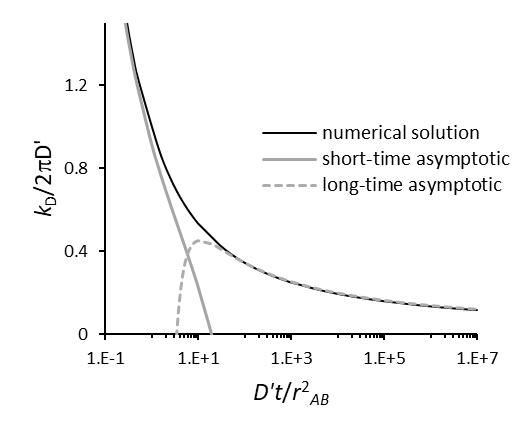
A general mass-acton applicability criterion for 2D can be formulated quantitatively by applying the Smoluchowski theory, which was extensively reviewed in the past, see, e.g., (Keizer, 1987); for a more recent overview, see (Yogurtcu and Johnson, 2015) and references therein. Using this approach, the kinetics of bimolecular binding in 2D differs significantly from that in three-dimensional (3D) spaces, for which the Smoluchowski model predicts mass-action kinetics for both reaction-limited and diffusion-limited modes. The purely diffusion limited rate in 3D (i.e. assuming infinitely fast initrinsic binding) may be characterized by the Smoluchowski reaction rate constant $k_{D}=4\pi r_{AB}D^{'}$, where $r_{AB}$ is the sum of effective radii of the binding partners, $r_{AB}=r_{A}+r_{B}$ , and $D^{'}$ is the sum of their diffusion coefficients, $D^{'}=D_{A}+D_{B}$. For the intermediate regime, characterized by finite $k_{0}$ and $k_{D}$ and termed “diffusion-influenced”, the theory also yields the mass-action kinetics (Collins and Kimball, 1949), with the rate constant

$k=\frac{k_{0}k_{D}}{k_{0}+k_{D}}$ .	Eq (S10)

In contrast, for the diffusion-limited case in 2D, the theory yields *time-dependent* rate coefficients $k_{D}\left( t \right)$ (Torney et al., 1983), i.e. mass-action kinetics do not apply to the diffusion-limited regime in 2D (the same is true for bimolecular binding in one dimension (Barzykin and Tachiya, 1993)). This is a consequence of the loss of dimensionality, which reduces the probability for a molecular pair to escape an encounter to zero, resulting in the formation of depletion zones.

In Figure S7, we determined the “universal” time dependence of the rate coefficient for the diffusion-limited binding in 2D, with $k_{D}$ scaled by $2\pi D^{'}$ and $t$ scaled by $\frac{r_{AB}^{2}}{D^{'}}$. (black curve). It was obtained by solving the Smoluchowski model numerically with *Virtual Cell* (VCell), a software system conataining numerical tools for solving models arising in
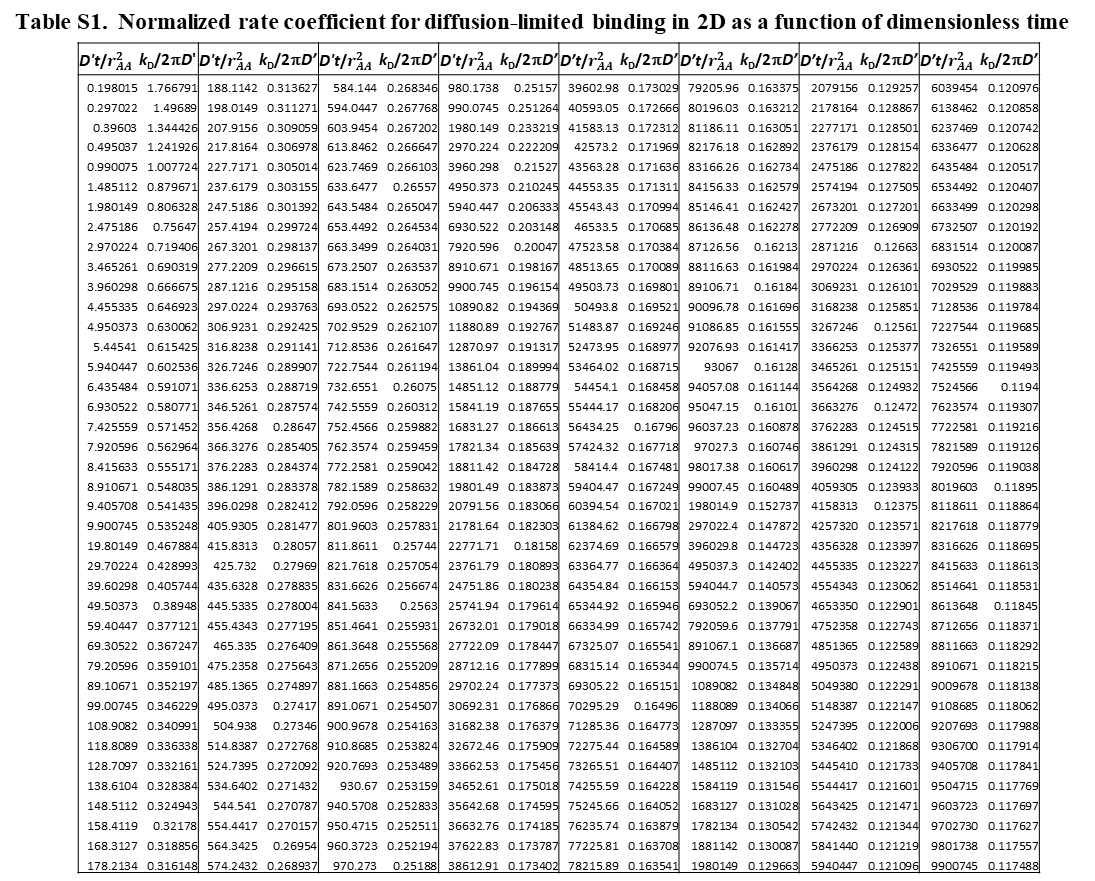
cell biology (Schaff, et al., 1997; Slepchenko et and Loew, 2010; Cowan, et al., 2012). The values of $k_{D}/2\pi D'$ as a function of $D^{'}t/r_{AB}^{2}$are shown in Table S1. The details for Smoluchowski model used to produce the solid black curve in Figure S7 and Table S1, are provided at the end of this Supporting Text.

**Figure S7. Time dependence of** $\boldsymbol{k}_{\boldsymbol{D}}$ **in 2D systems.**

Dimensionless rate constant$\kappa_{D}=$ $\frac{k_{D}}{2\pi}$, obtained numerically as a function of $\tau=\frac{D^{'}t}{r_{AB}^{2}}$ (black solid curve), is compared with analytical short-time and long-time asymptotic solutions (Torney, McConnell, 1983; Barzykin and Tachiya, 1993): $\kappa_{D}=\frac{1}{\sqrt{\pi\tau}}+\frac{1}{2}-\frac{1}{4}\sqrt{\frac{\tau}{\pi}}$ +… (grey solid curve) and $\kappa_{D}=2\left( \xi-\gamma\xi^{2}+\left( \frac{\pi^{2}}{6}-\gamma^{2} \right)\xi^{3}+\ldots\right),$ where $\xi=\frac{1}{\ln\left( 4\tau\right)-2\gamma}$ and Euler’s constant $\gamma=0.5772157$… . (grey dashed curve).

We now turn to the intermediate regime with finite intrinsic rate constants $k_{0}$. According to its analytical asymptotics, the decreasing function $k_{D}\left( t \right)$ approaches zero as $t\to\infty$ and diverges to infinity when $t\to0$. This suggests that in the intermediate regime with finite intrinsic rate constants $k_{0}$, the bimolecular binding in 2D begins as reaction-limited and thus initially it is well approximated by mass-action kinetics, but eventually it crosses over to the diffusion-limited mode. Then the question is, whether the mass-action kinetics has remained a good approximation up to the time of the reaction’s essential completion.

Suppose we would like to evaluate whether mass-action kinetics is a good approximation up to the time $t_{c}$corresponding to a completion level of the reaction, $p=$ $1-\frac{\sigma\left( t_{c} \right)}{\sigma_{0}}$ . If this is the case, Eq (S9) holds with ${k\approx k}_{0}$, and $t_{c}\cong\frac{1}{2k_{0}\sigma_{0}}\frac{p}{\left( 1-p \right)}.$ For a given $t_{c}$, the deviation of the kinetics of bimolecular binding in 2D from the mass-action rate law can be assessed by the following relative measure,

$\delta=\left( 1+\frac{1}{k_{0}t_{c}}\int_{0}^{\infty} k_{D}\left( t \right)\exp\left( -\frac{t}{t_{c}} \right)dt \right)^{-1}.$	Eq (S11)

We derived Eq (S11) from the general relation between the pair survival probabilities in the diffusion-limited and intermediate regimes (Pedersen, 1980), which holds for the systems of all dimensions (Tachiya, 1983 (Appendix)). This property, originating from linearity of the Smoluchowski model and the fact that the two regimes differ only by conditions at the reactive boundary and not by the equation itself, allows one to connect the kinetic coefficients of the two regimes (Szabo, 1989),

$\hat{k}\left( s \right)=k_{0}\hat{k}_{D}(s)/(k_{0}+s\hat{k}_{D}(s))$,	Eq (S12)

where $\hat{k}_{D}\left( s \right)=\int_{0}^{\infty} k_{D}\left( t \right)e^{-st}dt$ is the Laplace transform of $k_{D}\left( t \right)$, and $\hat{k}\left( s \right)=\int_{0}^{\infty} k\left( t \right)e^{-st}dt$ is the Laplace transform of the rate coefficient in the intermediate regime with a finite $k_{0}$; the Laplace variable $s$ has units of the inverse time. Note that in 3D, where both $k$ and $k_{D}$ are constants, Eq (S12) reduces to Eq (S10).

In lower dimensions, the meaning of $s\hat{k}_{D}\left( s \right)=\frac{1}{s^{-1}}\int_{0}^{\infty} k_{D}\left( t \right)exp(-\frac{t}{s^{-1}})dt$ is that of a weighted average of $k_{D}\left( t \right)$over $t\sim s^{-1}$, given that the main contributions to the integral come from the times $t$ that do not significantly exceed $s^{-1}$; we therefore denote $s\hat{k}_{D}\left( s \right)$ as $\bar{k}_{D}(s^{-1})$:

$\bar{k}_{D}(s^{-1})=\frac{1}{s^{-1}}\int_{0}^{\infty} k_{D}\left( t \right)exp(-\frac{t}{s^{-1}})dt$	Eq (S13)

While Eq (S12) holds for arbitrary $s$, we are interested in $s^{-1}=t_{c}\cong\frac{1}{2k_{0}\sigma_{0}}\frac{p}{\left( 1-p \right)}$, so from Eq (S12),

$\bar{k}\left( t_{c} \right)=k_{0}\bar{k}_{D}(t_{c})/(k_{0}+\bar{k}_{D}(t_{c}))$.	Eq (S14)

We now define a measure of relative deviation from mass-acton kinetics as $\delta=\bar{k}\left( t_{c} \right)/\bar{k}_{D}(t_{c})$; then from Eq (S14),

$\delta=\bar{k}\left( t_{c} \right)/\bar{k}_{D}(t_{c})={(1+\bar{k}_{D}(t_{c})/k_{0})}^{-1}$,

which, together with Eq (S13), brings us to Eq (S11).

For a given time to completion, *t_c_*, the closer *δ* is to 1, the more the rate will deviate from mass action and the ability to reliably use a time-independent rate constant. To apply Eq (S11) to dimerization of monomers with short tethers, we interpret the intrinsic binding rate constant $k_{0}$ as $k_{on}^{\left( h \right)}$, the effective reaction radius $r_{AA}$ as $2h$, and the coefficient of relative diffusion $D^{'}$ as $2D_{mem}$. Utilizing the tabulated values of $k_{D}/2\pi D^{'}$ as a function of $D^{'}t/r_{AA}^{2}$ (Table S1) to evaluate the integral in Eq (S11) numerically, one can compute $\delta$ for any combination of the equivalent 2D on-rate constant of association of binding sites ($k_{on}^{\left( h \right)}$), tether legnth ($h$), anchor diffusion coefficient ($D_{mem}$), initial monomer concentration ($\sigma_{0}$ ) and level of reaction completion ($p$). Table S2 shows values of $\delta$ computed for $h=$ 5.5 nm, $D_{mem}=$0.05 μm^2^/s, $p=$ 0.9, and selected values of $k_{on}^{\left( h \right)}$ and $\sigma_{0}$.

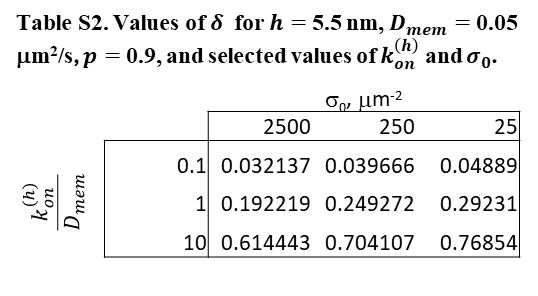

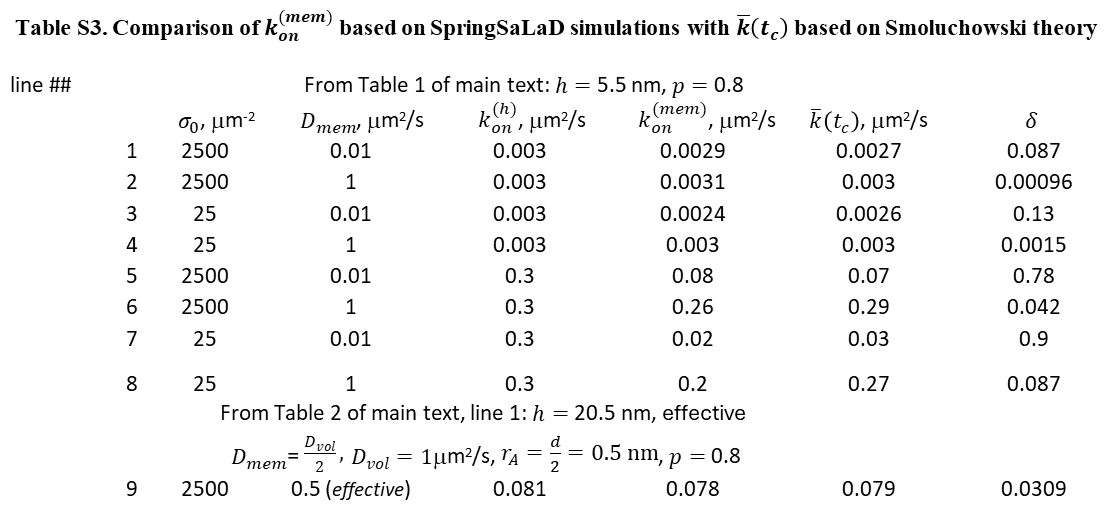
We also computed average values of the binding rate coefficient in the intermediate regime $\bar{k}\left( t_{c} \right)$ (Eq (S14)) using $p=$ 0.8 and the parameter values of Table 1 of the main text to compare them with the corresponding values of $k_{on}^{\left( mem \right)}$. The latter are also averages of $k(t)$, as they were obtained by fitting Eq (S9) to SpringSaLaD simulations run to the 80% completion of dimerization. The comparison showed reasonably good agreement (rows ##1-8 of Table S3), giving credence to both the simulations and theory. Also, shown in Table S3 are the corresponding values of *δ*. As illustrated in Figure 2 of the main text, the parameters associated with row #5 of Table 1 produce kinetics in SpringSaLaD that do not fit well to the mass action rate law; the corresponding value of *δ*, 0.78 (row 5 of Table S3), is indeed close to 1. As a rule of thumb, we propose a value of *δ* < 0.3 to allow for the application of 2D mass action as a reasonable approximate rate law.

In the case of long tethers, $h>\left( \pi\sigma_{0} \right)^{-\frac{1}{2}}$, exemplified by the first row of Table 2 of the main text, geometric constraints imposed by tethers and slow anchor diffusion are less limiting in two aspects. First, multiple binding sites may now interact under the hemisphere of radius $h$. As a result, the tether length no longer represents the effective reaction radius, which is now determined by the radii of the binding sites $d/2$. Second, the main mixing mechanism is now volumetric diffusion. Because the area of the hemisphere is twice the area of its projection on the membrane, and both are covered by diffusion within the same time, the corresponding effective 2D diffusivity is $D_{vol}/2$. Thus, to compute $\delta$ and $\bar{k}\left( t_{c} \right)$ for the case of long tethers, we should use Eqs (S11) and (S14) with $r_{AA}=d$ and $D^{'}=$ $D_{vol}$.

The value of $\bar{k}\left( t_{c} \right)$ in row #9 of Table S3, obtained with the same parameters as in the first row of Table 2 of the main text, is in good agreement with the $k_{on}^{\left( mem \right)}$, obtained by fitting Eq (S9) results from the SpringSaLaD simulations, and it is characterized by low $\delta$, i.e. the kinetics of this example is well approximated by the mass-action rate law.

Note that the condition $h>\left( \pi\sigma\right)^{-\frac{1}{2}}$ may not hold for all levels of completion $p$. Indeed, the right-hand side of the inequality increases as $\sigma$ drops. The effective reach of the binding side, represented by the left-hand side, would also increase due to the anchor’s diffusion. Hence, one can estimate the maximum completion level $p_{max}$ , at which the long-tether condition holds, by solving the equation, $h+2{(D_{mem}t(p))}^{1/2}=\left( \pi(1-p )\sigma_{0} \right)^{-1/2}$.The dependence $t(p)$ is the inverse of the time dependence of dimer density, which can be retrieved from the SpringSaLaD simulation results. One can also solve the above equation directly assuming applicability of Eq (S9), from which $t\left( p \right)=\frac{1}{2k_{on}^{\left( h \right)} \sigma_{0}}\frac{p}{\left( 1-p \right)}$. Substituting this in the equaiton above yields $h\sqrt{\pi\left( 1-p \right)\sigma_{0}}+\sqrt{\frac{2\pi D_{mem}}{k_{on}^{\left( h \right)}}}p=1$, which is solved exactly. For the parameters of the example in row #9 of Table S3: $h=0.02$ μm, $\sigma_{0}=$ 2500 μm^2^, $D_{mem}=$ 0.01 μm^2^/s, and $k_{on}^{\left( h \right)}=0.081$ μm^2^/s, we find $p_{max}=0.995$. Thus, the completion level $p=$0.8, used for this example in SpringSaLaD and in estimating the corresponding $\bar{k}\left( t_{c} \right)$, satisfies the long-tether condition.

The mass-action rate law for reversible homodimerization is $\partial_{t}\sigma=-2k\sigma^{2}+kK(\sigma_{0}-\sigma)$, where $K$ is the equilibrium dissociation constant. Its exact solution $x(t)\equiv\frac{\sigma(t)}{\sigma_{0}}$ satisfies the following relation,

$\frac{x(t)-x_{1}}{x\left( t \right)-x_{2}}=\frac{1-x_{1}}{1-x_{2}}\mathrm{ex}p \left( -2k\sigma_{0}\sqrt{x_{1}-x_{2}}t \right)$,	Eq (S15)

where $x_{1}$ is the normalized steady state, $x_{1}=\frac{\sigma_{stedy state}}{\sigma_{0}}=\frac{1}{2}(\sqrt{\alpha^{2}+4\alpha}-\alpha)$ and $x_{2}=-\frac{1}{2}(\sqrt{\alpha^{2}+4\alpha}+\alpha)$, where $\alpha=K/2\sigma_{0}$.

In the reversible case, the level of completion $p$ reflects how close the reaction is to its steady state, i.e. $1-p=$ $x\left( t \right)-x_{1}.$ Then from Eq (S15), $\frac{1-p}{1-p+x_{1}-x_{2}}=\frac{1-x_{1}}{1-x_{2}}e^{-2k\sigma_{0}\sqrt{x_{1}-x_{2}}t}$, and the corresponding time to completion $t_{c}$ in this case is

$$t_{c}\cong\frac{1}{2k_{0}\sigma_{0}\sqrt{x_{1}-x_{2}}}\ln\left( \frac{(1-p+x_{1}-x_{2})(1-x_{1})}{\left( 1-p \right)(1-x_{2})} \right).$$

*Computation details related to Figure S7 and Table S1*

The Smoluchowski model defines the binding rate coefficient $k(t)$ in terms of pair survival probability $u(r,t)$, the probability that an isolated pair of binding partners, initially separated by distance $r$, does not react by time $t$. Here and below, we assume uniformity of space, system axial symmetry, and that irreversible binding occurs on encounter. Without loss of generality, one of the binding partners (molecule A) can be fixed at the origin and assigned the effective reaction radius, $r_{AB}=r_{A}+r_{B}$, whereas the center of the other binding partner (molecule B) diffuses with the effective diffusion coefficient $D^{'}=D_{A}+D_{B}$ in the domain $r\in[r_{AB}, \infty)$. The survival probability $u$ is governed by the equation (see, e.g., Barzykin and Tachiya, 1993), $u_{t}=D'\Delta u$, where $\Delta$ is the 2D diffusion operator, $\Delta=\frac{1}{r}\partial_{r}(r\partial_{r} )$. The governing equation is solved with the flux boundary condition, $2\pi r_{AB}\partial_{r}u\left( r_{AB},t \right)=k_{0}u\left( r_{AB},t \right)$, where $k_{0}$ is the rate constant of intrinsic binding, the initial condition $u\left( r,0 \right)=1$, and the boundary condition at $r_{AB}\to\infty$ $u\left( \infty,t \right)=1$. The reacton rate coefficient $k\left( t \right)$ is then defined as $k(t)=2\pi r_{AB}\partial_{r}u\left( r_{AB},t \right)$. For diffusion-limited reactions, $k_{0}\to\infty$, so the boundary condition at $r=r_{AB}$ becomes $u\left( r_{AB},t \right)=0$. Thus, $k_{D}(t)=$ $2\pi r_{AB}\partial_{r}u\left( r_{AB},t \right)$ for the reactive boundary condition $u\left( r_{AB},t \right)=0$.

To solve this model with VCell, the governing equation must be rewritten in a 1D ‘Cartesian’ diffusion-advection form: using the change of variables, $u=U/r$, the equation becomes $U_{t}=D^{'}\partial_{rr}^{2}U-\partial_{r}(vU)$, with the ‘advection velocity’ $v=D^{'}/r$. To avoid numerical spatial differentiation, which is lower order of accuracy than numerical integration, we solved the model both outside and inside the reaction boundary. Outside the reactive boundary, $r\in[r_{AB}, r_{max}]$ , we solve for $U$ using the flux density $f_{U}\left( r_{AB},t \right)=-k_{0}U\left( r_{AB},t \right)$at the reactive boundary with sufficiently large $k_{0}$, the initial condition $U(r,0)=r$, and the boundary condition $f_{U}\left( r_{max},t \right)=0$ at $r=r_{max}$. Inside the reactive boundary, $r\in[0,r_{AB})$, we integrate the flux crossing the reactive boundary over time, $I\left( t \right)=\int_{0}^{t} k_{D}(t')dt'$ . For this, we solve there the 1D diffusion equation, $u_{t}=D\partial_{rr}^{2}u,$ with sufficiently high diffusivity $D$, so that $u(r,t)$ is equal to its average $\bar{u}(t)$ with high precision. The equation was solved with the flux boundary conditions, $f_{u}\left( r_{AB},t \right)=f_{U}\left( r_{AB},t \right)/r$ and $f_{u}\left( 0,t \right)=0$, and the intial condition $u(r,0)=0.$ Thus, $\bar{u}(t)r_{AB}$ yields the integrated flux density and the integrated flux $I\left( t \right)$ is given by $2\pi r_{AB}^{2}\bar{u}(t)$. Finally we determine $k_{D}(t)$ by numerical time differentiation of $\bar{u}(t)$, $k_{D}\left( t \right)=\frac{dI}{dt}=2\pi r_{AB}^{2}\bar{u}_{t}.$ The computations were performed using VCell’s fully-implicit finite-volume solver (Slepchenko et al, 2018; Resasco, et al., 2012) for $r_{AB}=0.001005$ μm, $r_{max}=10$ μm, $D^{'}=0.01$ μm^2^/s, $D=1$ μm^2^/s. Results obtained with the increasing $k_{0}$ and decreasing space discretization parameter $\Delta r$, showed that the relative numerical errors of the solution with $k_{0}={10}^{6}$ μm^2^/s and $\Delta r={10}^{-5}$ μm were within few percent. Comparison against the short-time and long-time analytical asymptotic solutions (Figure S7) indicated relative errors in the 2-4% range. We nondimensionalized the solution obtained with $k_{0}={10}^{6}$ μm^2^/s and $\Delta r={10}^{-5}$ μm to obtain the universal dependence of $k_{D}/2\pi D'$ on normalized time $\tau=D^{'}t/r_{AB}^{2}$ (Table S1). The VCell implementation of the model can be found in the public VCell MathModel database under username ‘boris’. Model name: Diffusion_limited_binding_2D’; simulations: Simulation3 and its copies.
